# Emergence of Emotion Selectivity in Deep Neural Networks Trained to Recognize Visual Objects

**DOI:** 10.1101/2023.04.16.537079

**Authors:** Peng Liu, Ke Bo, Mingzhou Ding, Ruogu Fang

## Abstract

Recent neuroimaging studies have shown that the visual cortex plays an important role in representing the affective significance of visual input. The origin of these affect-specific visual representations is debated: they are intrinsic to the visual system versus they arise through reentry from frontal emotion processing structures such as the amygdala. We examined this problem by combining convolutional neural network (CNN) models of the human ventral visual cortex pre-trained on ImageNet with two datasets of affective images. Our results show that (1) in all layers of the CNN models, there were artificial neurons that responded consistently and selectively to neutral, pleasant, or unpleasant images and (2) lesioning these neurons by setting their output to 0 or enhancing these neurons by increasing their gain led to decreased or increased emotion recognition performance respectively. These results support the idea that the visual system may have the intrinsic ability to represent the affective significance of visual input and suggest that CNNs offer a fruitful platform for testing neuroscientific theories.

**Author Summary:** The present study shows that emotion selectivity can emerge in deep neural networks trained to recognize visual objects and the existence of the emotion-selective neurons underlies the ability of the network to recognize the emotional qualities in visual images. Obtained using two affective datasets (IAPS and NAPS) and replicated on two CNNs (VGG-16 and AlexNet), these results support the idea that the visual system may have an intrinsic ability to represent the motivational significance of sensory input and CNNs are a valuable platform for testing neuroscience ideas in a way that is not practical in empirical studies.

## Introduction

Human emotions are complex and multifaceted and under the influence of many factors, including individual differences, cultural backgrounds, and the context in which the emotion is experienced (*1–5*). Still, a large number of people, across different cultures, different levels of education, and different socioeconomic backgrounds, experience similar feelings when viewing images of varying affective content (*6–9*). What fundamental principles in the functions of the human visual system underlie such universality requires elucidation.

Previous studies of emotion perception have primarily relied on empirical cognitive experiments (*10–12*). Some of them have focused on capturing human behavioral valence or arousal judgment on affective images (*13–16*), while others have recorded brain activities to look for neural correlates of affective stimuli processing (*17–21*). Despite decades of effort, how the brain transforms visual stimuli into subjective emotion judgments (e.g., happy, neutral, or unhappy) remains not well understood. The advent of machine learning especially artificial neural networks (ANNs) opens the possibility of addressing this problem using a modeling approach.

Artificial neural networks can project visual images to a feature space in which the activation patterns of hidden layers are the features used for object classification and recognition. One type of artificial neural network, convolutional neural networks (CNNs), owing to their hierarchical organization resembling that of the visual system, are increasingly used as models of visual processing in the primate brain (*22–26*). CNNs trained to recognize visual objects can achieve performance levels rivaling or even exceeding that of humans. Interestingly, CNNs trained on images from such databases as ImageNet (*27*) are found to demonstrate neural selectivity for a variety of stimuli that are not included in the training data. For instance, (*28*) showed that neurons in a CNN trained on ImageNet became selective for numbers without having been trained on any “number” datasets. Similarly, (*29*) demonstrated that a CNN, when trained on non-face objects, can develop a recognition performance for faces that significantly exceeds chance levels. These instances demonstrate that CNNs may possess recognition capabilities beyond the primary task they are trained on.

The role of the visual cortex in visual emotion processing is debated (*30*, *31*). (*32*) argued that emotion representation is an intrinsic property of the visual cortex. They used a CNN pre-trained on ImageNet to show that the model can accurately predict the emotion categories of affective images. (*20*), on the other hand, showed that the affective representations found in the visual cortex during affective scene processing might arise as the result of reentry from anterior emotion-modulating structures such as the amygdala. The goal of this study is to further examine this question using CNN models.

CNN models are well suited for addressing questions related to the human visual system. Among the many well-established CNN models, VGG-16 (*33*) has an intermediate level of complexity and is shown to have superior object recognition performance (*34*). Using VGG-16, recent cognitive neuroscience studies have explored how encoding and decoding of sensory information are hierarchically processed in the brain (*23*, *35*, *36*). (*23*) used VGG-16 to quantitatively demonstrate an explicit gradient of feature complexity encoded in the ventral visual pathway. (*35*) used VGG-16 to model the visual cortical activity of human participants viewing images of objects and demonstrated that activities in different layers of the model highly correlate with brain activities in different visual areas. (*36*) investigated qualitative similarities and differences between VGG-16 and other feed-forward CNNs in the representation of the visual object and showed these CNNs exhibit multiple perceptual and neural phenomena such as the Thatcher effect (*37*) and Weber’s law (*38*).

In this study, we mainly focused on VGG-16 pre-trained on ImageNet as the model of the human visual system and used AlexNet (*39*), which is another well-established CNN model of visual processing, to test whether the results can be replicated. Using two well-established affective image datasets: International Affective Picture System (IAPS) (*15*) and Nencki Affective Picture System (NAPS) (*16*), we examined whether emotion selectivity can spontaneously emerge in such systems and whether such emotion selectivity has functional significance. For each filter within a layer of the model, the emotional selectivity for the resulting feature map was established by first computing neural responses to three broad classes of images: pleasant, neutral, and unpleasant (tuning curves) at the level of each unit and then averaging these responses across all the units within the feature map. A feature map, also referred to as a neuron in what follows, is considered selective for a particular emotion if its tuning responses are robust and exhibit the strongest responses to images of that category from both datasets. To test whether these emotion-selective neurons have a functional role, we replaced the last 1000-unit object-recognition layer of the VGG-16 with a two-unit emotion-recognition layer and trained the connections to this layer to decode pleasant versus non-pleasant, neutral vs. non-neutral, and unpleasant vs. non-unpleasant images. Two neural manipulations were carried out: lesion and feature attention enhancements. Lesioning the neurons selective for a specific emotion is expected to degrade the network’s performance in recognizing that emotion, whereas applying attention enhancement to the neurons selective for the emotion is expected to increase the network’s performance in recognizing that emotion.

## Results

We tested whether emotion selectivity can naturally arise in a CNN model trained to recognize visual objects. VGG-16 pre-trained on ImageNet data (*27*) was used for this purpose (see Figure 1). Filters/channels within a layer were referred to as neurons and responses from the units within the feature maps were averaged and treated as neuronal responses. Selectivity for pleasant, neutral, and unpleasant emotions was defined for each neuron based on its response profiles to images from two affective picture sets (IAPS and NAPS). The functional significance of these neurons was then assessed using lesion and attention enhancement methods.

**Fig. 1.**
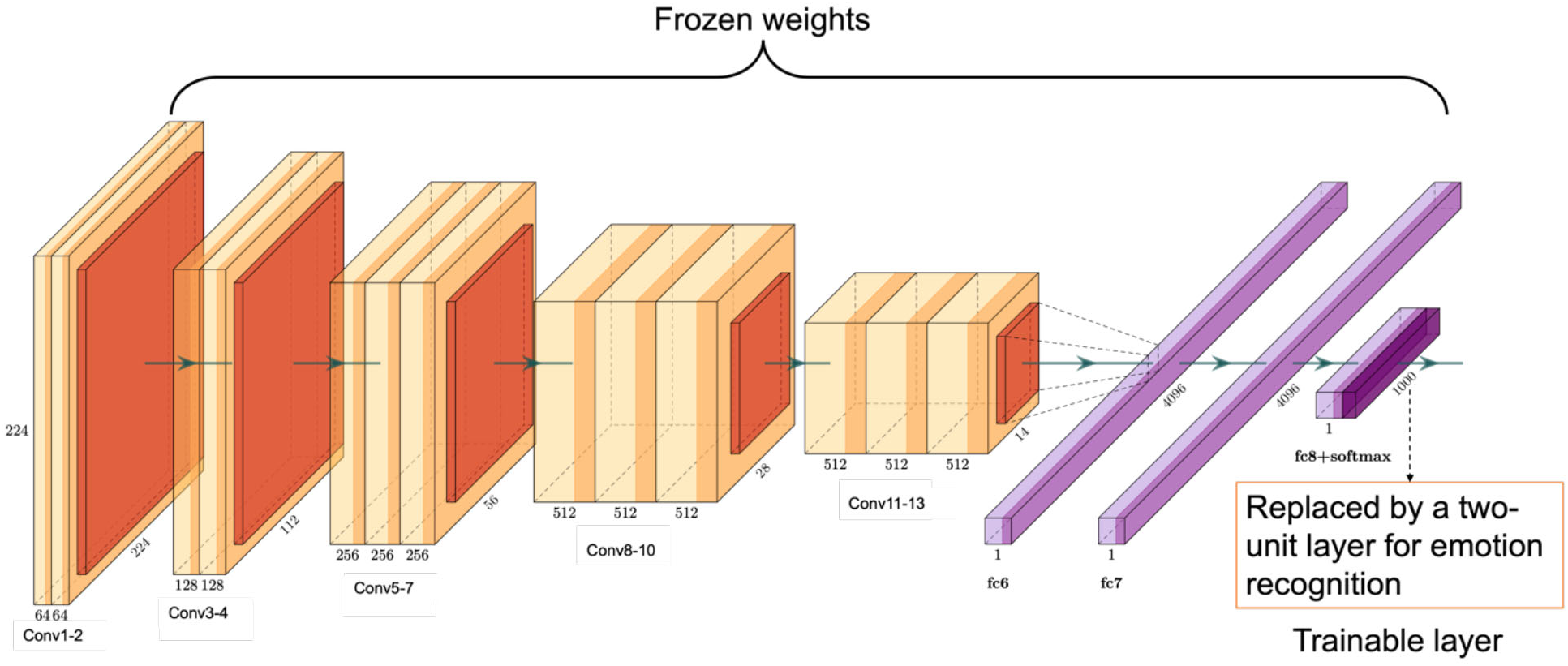
The architecture of the VGG-16 model. We used the VGG-16 pre-trained on ImageNet to model the visual system. VGG-16 has 13 convolutional layers and three fully connected (FC) layers. Each convolutional layer (light yellow color) is followed by a ReLU activation layer (yellow color) and a max-pooling layer (red color). Each FC layer (light purple color) is followed by a ReLU layer (purple color). The last FC layer is followed by a ReLU and a SoftMax layer (dark purple color). In the original VGG-16, the last layer was used to recognize 1000 different objects. In our model it was replaced by a two-unit layer whose connections to the preceding layer were trained to recognize different emotions: (1) pleasant vs. non-pleasant; (2) neutral vs. non-neutral; (3) unpleasant vs. non-unpleasant. Affective images in grayscale from two datasets (IAPS and NAPS) were presented to the model to define the emotion-selectivity of neurons in the convolutional layers. Lesion and attention enhancement were applied to assess these neurons’ functional significance.

### Neuronal responses to emotional images in different convolutional layers

The tuning curve for a neuron is defined as the normalized mean response (tuning value) to pleasant, neutral, and unpleasant images in a given dataset plotted as a function of the emotion category. The maximum of the tuning curve indicates the neuron’s preferred emotion category for that picture set. Figure 2A (top) shows the tuning curves of three neurons from the Convolutional Layer 3 (an early layer) for both IAPS and NAPS datasets. According to the definition above, these neurons are selective for pleasant, neutral, and unpleasant categories, respectively. For the top 100 images from IAPS and NAPS that elicited the strongest response in these neurons, Figure 2A (bottom) shows the valence distribution of these images. As can be seen, for these early layer neurons, while the pleasant neuron is more activated by images with high valence ratings (pleasant), for the neutral and unpleasant neurons, the patterns are less clear. For the neurons in Convolutional Layer 6 (a middle layer), however, as shown in Figure 2B, their emotion selectivity and the category of images they prefer show greater agreement. Namely, the pleasant neuron prefers predominately images with high valence (pleasant), the neural neuron prefers predominately images with intermediate valence (neutral), and the unpleasant neuron prefers predominately images with low valence (unpleasant). The results for the three neurons from Convolutional Layer 13 (a deep layer) are similar to those from Layer 6; see Figure 2C.

**Fig. 2.**
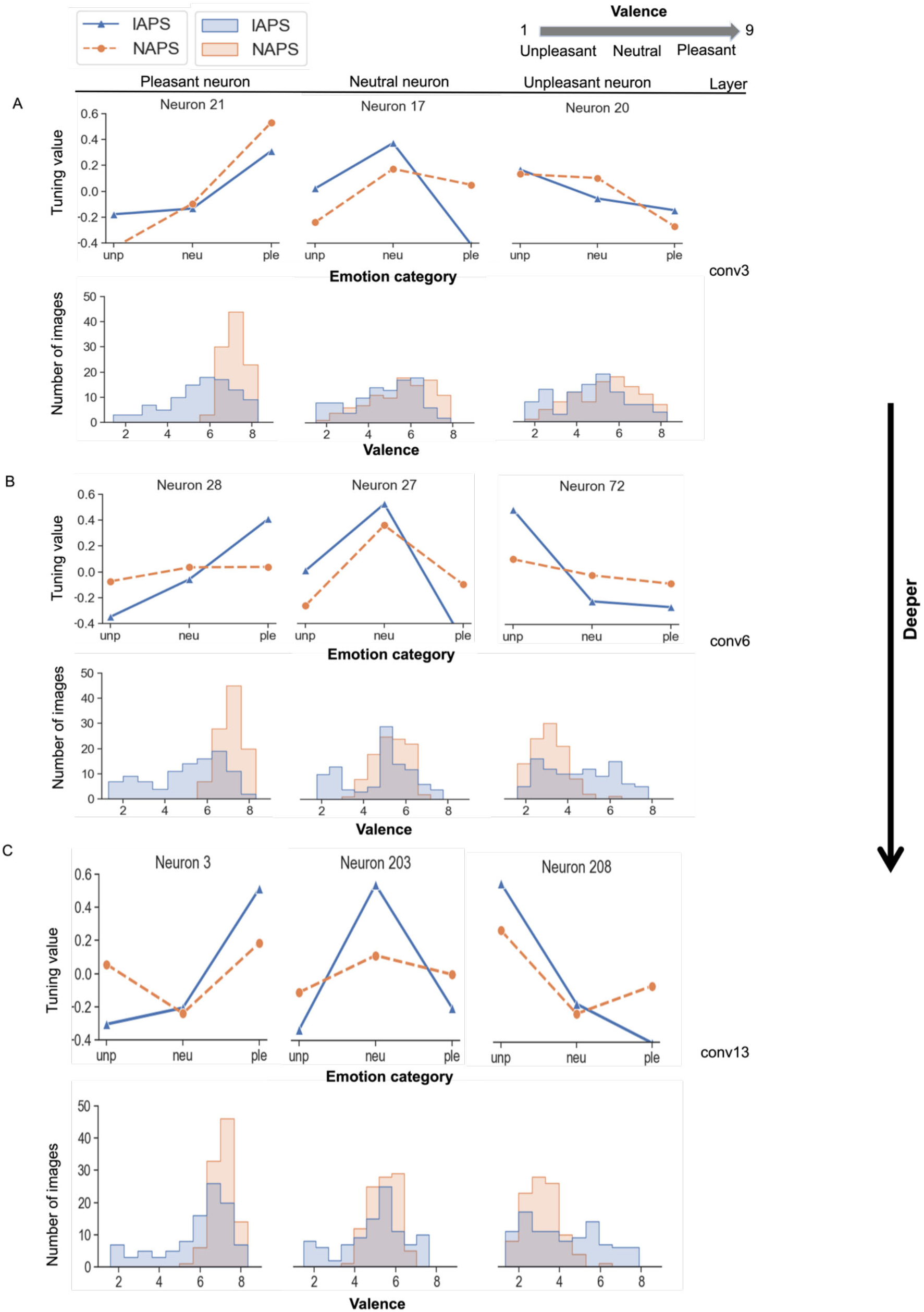
Tuning curves and emotion selectivity. (**A**-**C**) Tuning curves of example neurons from different convolutional layers (top panel) along with the valence distribution of the top 100 images that elicited the strongest responses for a given neuron.

### Emotion selectivity in different convolutional layers

Whereas tuning value and tuning curve characterize a neuron’s response to images from different emotion categories, the selectivity index (SI), which highlights the difference between responses to different emotion categories of images, is a better index for defining emotion selectivity. As shown in Figure 3A, emotion selectivity became stronger as one ascended the layers from early to deep, an effect that is especially noticeable for the IAPS datasets, supporting the notion that emotion differentiability increases as we go from earlier to deeper layers. In light of the computational principle that earlier layer neurons encode lower-level stimulus properties (e.g., shapes and edges) and deeper layer neurons encode higher-level properties such as semantic meaning (e.g., object identities) (*40–42*), the results in Figure 3A as well as Figure 2 suggest that from earlier to deeper layers, emotion as a higher level cognitive construct becomes progressively better defined and better differentiated.

**Fig. 3.**
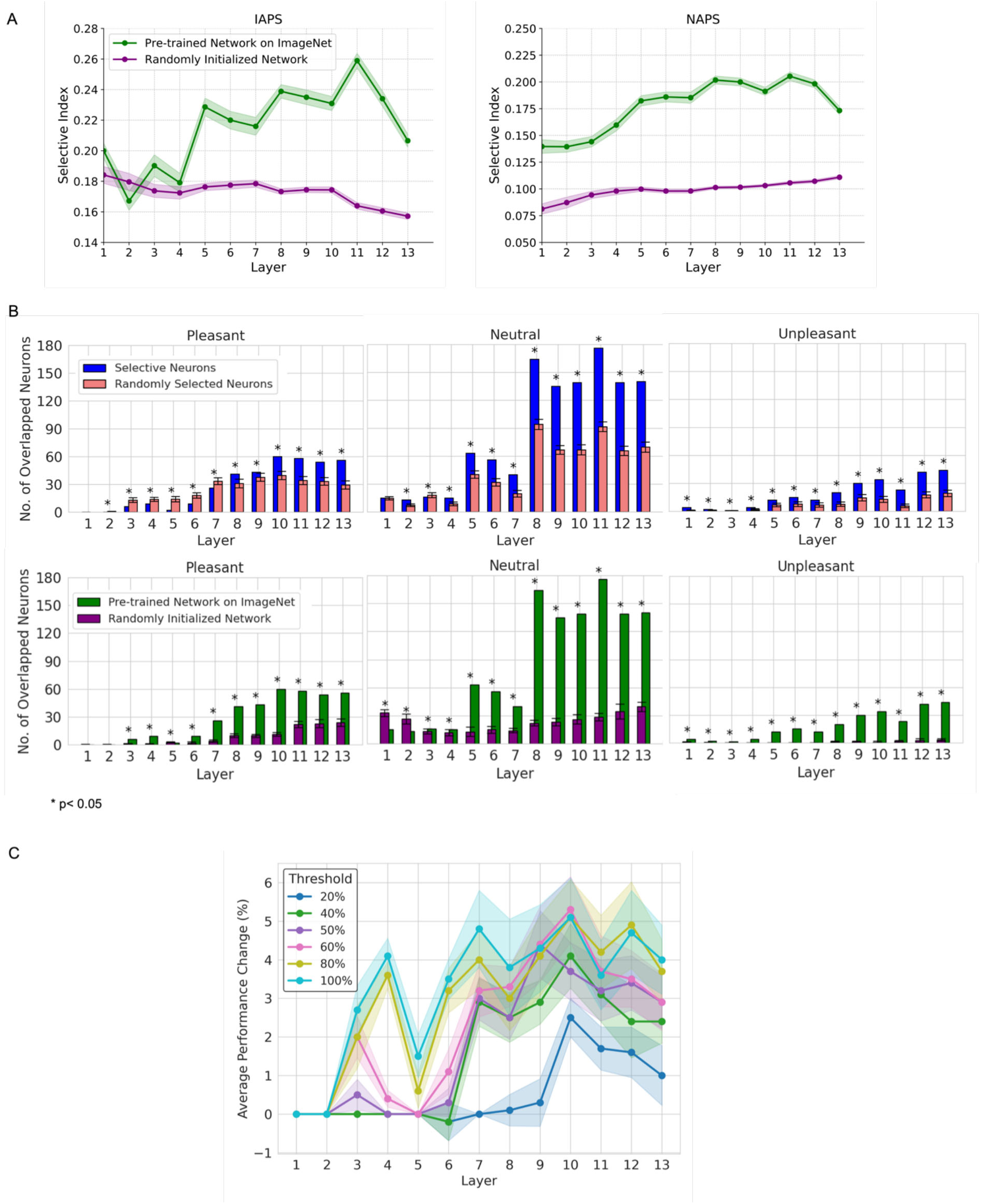
Emotion selectivity and its generalizability. (A) Emotion selectivity as a function of layer for IAPS and NAPS. (B-top) Number of neurons determined to be selective for a given emotion for both IAPS and NAPS datasets compared with the number of neurons in the overlap of two random sets of neurons. (B-bottom) The number of neurons determined to be selective for a given emotion for both IAPS and NAPS datasets in VGG-16 pretrained on ImageNet and with randomly initialized weights. (C) Removing successively larger percentages of neurons with small SI values and comparing the performance of attention-enhancing the remaining neurons yielded a threshold of 80% for determining emotion selectivity.

To examine the role of the training to recognize objects in the foregoing observations, we performed the same analysis in a VGG-16 with randomly initialized weights (i.e., not trained to recognize objects). As seen in Figure 3A, emotion selectivity is generally low as evaluated by both datasets, and there is no clear layer-dependence in emotion selectivity, suggesting that the increased ability to represent and differentiate emotion in deeper network layers of the pre-trained VGG-16 is an ability acquired through the training for object recognition.

### Generalizability of emotion-selective neurons

Figure 2 shows that a neuron can be tuned for the same emotion for both IAPS and NAPS datasets. A natural question is whether such neurons arise as the result of random chance or as an emergent property of the trained network. Further, based on the value of SI, all neurons are selectivity for one emotion or the other. Small SIs are likely subject to the influence of chance, and as such, neurons with small SIs should be removed from further consideration. How to determine the threshold for removal?

We performed two analyses to address the two questions. First, we rank-ordered neurons according to their SI values, removed certain percentages of neurons with small SI values, and attention-enhanced the remaining neurons (see next subsection) and observed the resulting performance improvement. The results in Figure 3C suggest that removing neurons whose SIs fell in the lower 20% (keeping 80%) is a reasonable threshold. Second, neurons determined to be emotion selective according to IAPS and that according to NAPS were subjected to an overlap analysis. Figure 3B (top) compares the number of neurons selective for the same emotion for both IAPS and NAPS datasets against the number of neurons to be expected from the overlap of two random sets of neurons. The former is consistently higher than the latter across all layers, with the effect becoming more prominent in deeper layers, suggesting that emotion selectivity generalizes across the two datasets and the generalizability is not due to chance.

What is the role of training to recognize visual objects in the generalizable emotion selectivity? To answer this question, we compared the number of emotion-selective neurons from the overlap analysis derived from pre-trained VGG-16 on ImageNet against that derived from randomly initialized VGG-16. Figure 3B (bottom) shows that for all emotion categories—pleasant, neutral, and unpleasant—the pre-trained network consistently demonstrated a higher number of emotion-selective neurons in the later layers, especially from Layer 5 onwards. These findings suggest that emotion selectivity is an emergent property as the result of a neural network undergoing training for object recognition.

### The functionality of emotion-selective neurons

To test whether emotion-selective neurons have a functional role, we followed (*43*) and replaced the last layer of the VGG-16, which originally contained 1,000 units for recognizing 1000 different types of objects, with a fully connected layer containing two units for recognizing two types of emotions. Three models were trained and tested for each of the two affective picture datasets: Model 1: pleasant versus non-pleasant, Model 2: neutral versus non-neutral, and Model 3: unpleasant versus non-unpleasant. Once these models were shown to have adequate emotion recognition performance (see Table 1), two neural manipulations were considered: feature attention enhancement and lesion. For feature attention enhancement (*44–46*), the gain of the neurons selective for a given emotion for both datasets is increased by increasing the slope of the ReLU activation function (see Methods) (*47–50*), whereas for lesion, the output of the neurons selective for a given emotion for both datasets was set to 0, which effectively removes the contribution of these neurons, i.e., they are lesioned. We hypothesized that (1) with attention enhancement, the network’s ability to recognize emotion is increased (2) with lesioning, the network’s ability to recognize emotion is decreased, and (3) such effects are not observed for modulating randomly selected neurons.

**Table 1.**
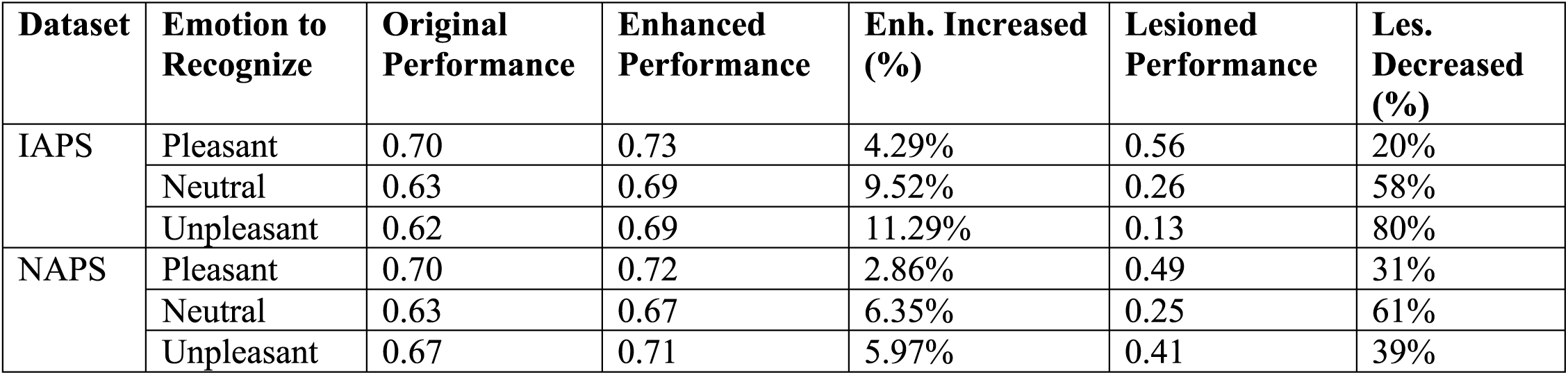
Original and Enhanced and Lesioned Performance (F1-score) in VGG-16. . The maximum performance changes for both enhancing and lesioning selective neurons across different layers are shown below.

#### Feature attention enhancement

For IAPS images, Figure 4A compares performance changes after enhancing the emotion-selective neurons as well as enhancing the same number of randomly sampled neurons; see also Table 1. The optimal tuning strength for which we achieved the best performance enhancement was chosen for each layer in the plot. As one can see, for pleasant versus non-pleasant, neutral versus non-neutral, and unpleasant versus non-unpleasant emotions, enhancing the gain of the neurons selective for a specific emotion can significantly improve the emotion recognition performance of the CNN model for that emotion. Moreover, deeper layer attention enhancement tends to yield greater performance improvements than earlier layer attention enhancement. Increasing the gain in randomly selected neurons, however, shows either a marginal performance improvement or a significant performance decline. The feature-attention performance of emotion-selective neurons over random neurons is highly statistically significant in the middle and deeper layers (p< 1.2e-02). Figure 4A (right) shows the performance changes across layers as the tuning strength varied from 0 to 5.

**Fig. 4.**
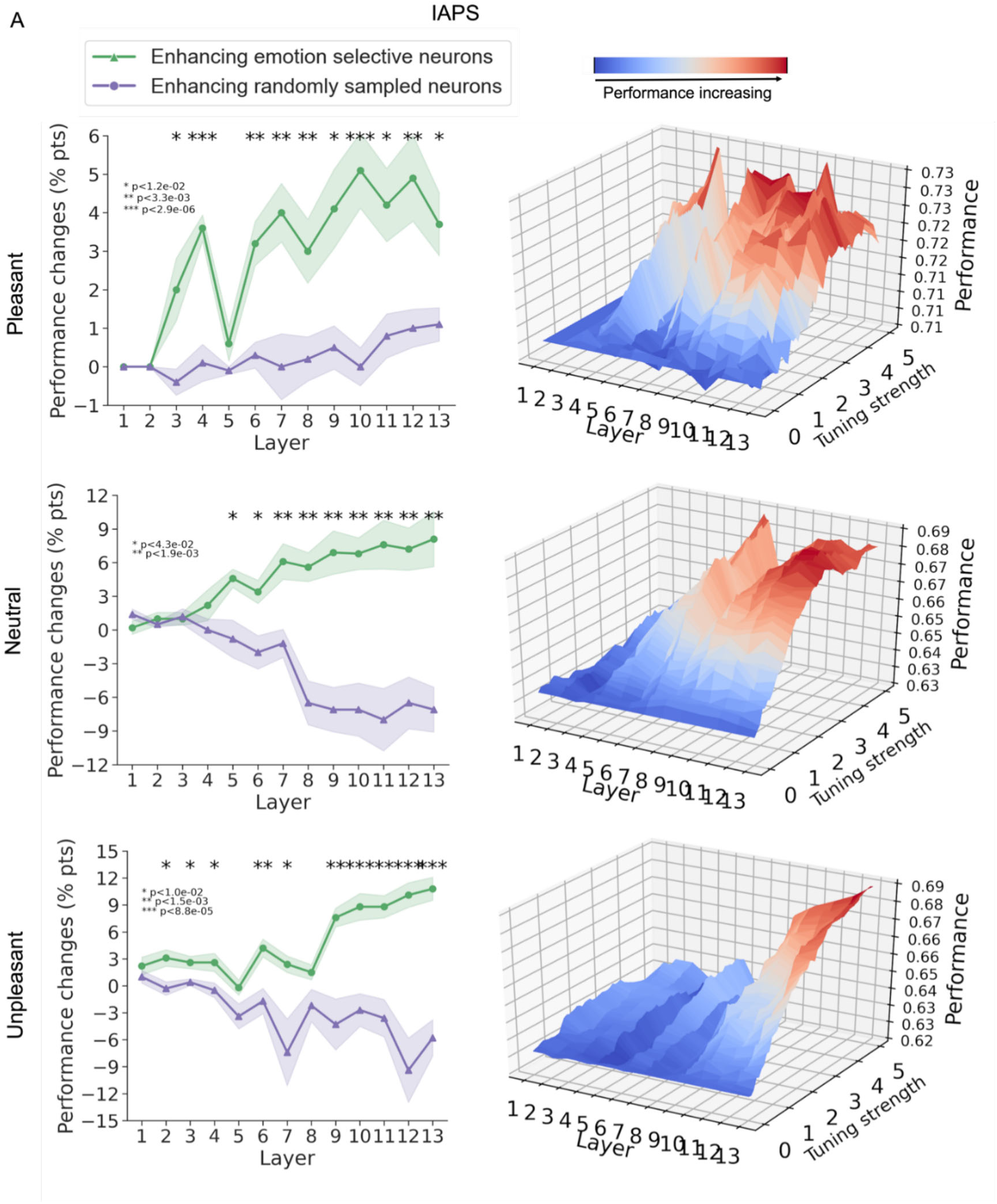
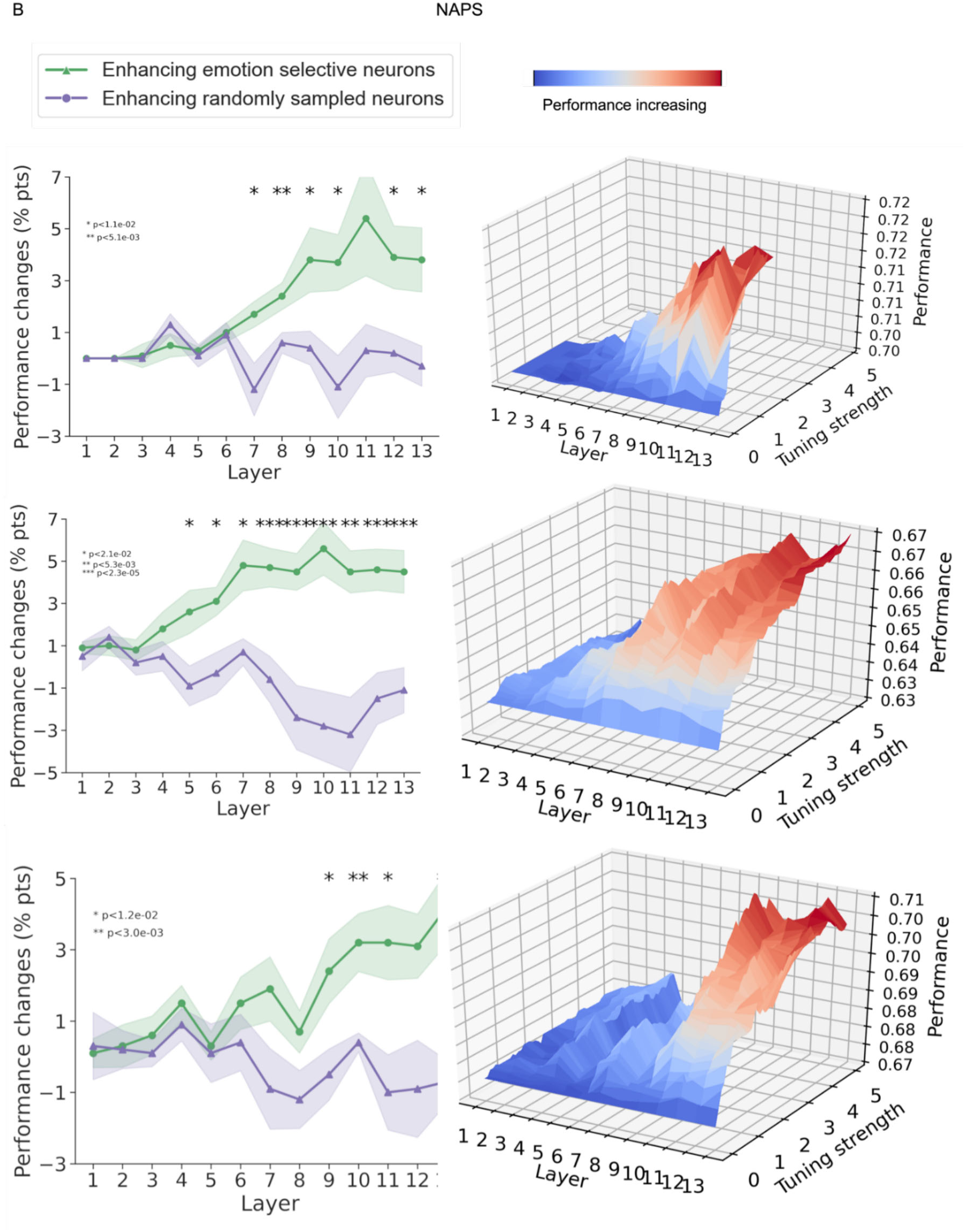
Effects of enhancing emotion-selective neurons and randomly selected neurons. (**A**) IAPS dataset. (**B**) NAPS dataset.

We carried out the same analysis for the NAPS dataset in Figure 4B. The results largely replicated that in Figure 4A for the IAPS dataset.

#### Lesion analysis

The functional importance of the emotion-selective neurons can be further assessed through lesion analysis (*51–54*). As shown in Figure 5 (see also Table 1), we compared the emotion recognition performance changes by setting the output from emotion-selective neurons to 0 as well as by setting the output of an equal number of randomly chosen neurons to 0. As can be seen, lesioning the emotion-selective neurons led to significant performance declines, especially for the deeper layers; the performance decline can be as high as 80%. In contrast, lesioning randomly selected neurons produces almost no performance changes. These results, replicated across both datasets, further support the hypothesis that emotion-selective neurons are important for emotion recognition, and the importance is higher in deeper layers than in earlier layers.

**Fig. 5.**
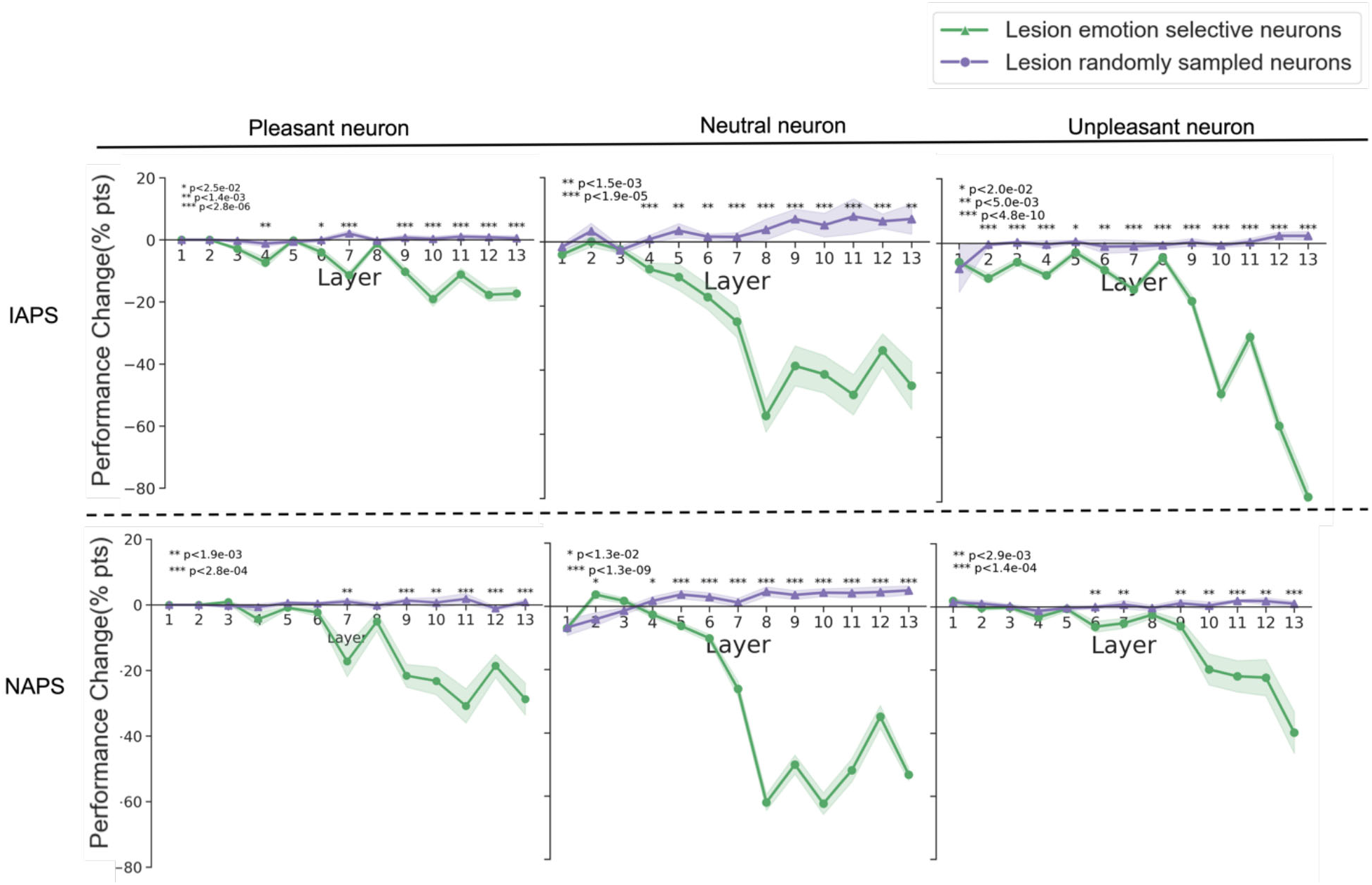
Lesion Analysis. Performance changes were compared between lesioning emotion-selective neurons and randomly selected neurons.

## Discussion

It has been argued that the human visual system has the intrinsic ability to recognize the motivational significance of environmental inputs (*55*). We examined this problem using convolutional neural networks (CNNs) as models of the human visual system (*56–61*). Selecting the VGG16 pre-trained on images from the ImageNet as our model (*62–64*) and using two sets of affective images (IAPS and NAPS) as test stimuli, we found the existence of emotion-selective neurons in all layers of the model even though the model has never been explicitly trained to recognize emotion. Additionally, emotion selectivity becomes stronger and more consistent in the deeper layers, in agreement with prior literature suggesting that the deeper layers of CNNs encode higher-level semantic information. For VGG-16 with randomly initialized weights (i.e., not trained to recognize objects), however, no such effects were observed, suggesting that emotion selectivity may be an emergent property through network training. Applying two manipulations: feature attention enhancement and lesion, we can show further that the emotion-selective neurons are functionally significant, specifically: (1) after increasing the gain of emotion-selective neurons (e.g., feature attention enhancement), the network’s performance in emotion recognition is enhanced relative to increasing the gain of randomly selected neurons and (2) in contrast, after lesioning the emotion-selective neurons, the network’s performance in emotion recognition is degraded relative to lesioning randomly selected neurons. These performance differences are stronger and more noticeable in deeper layers than in earlier layers. In the Supplementary Materials, we reported similar findings on the AlexNet, which is a simpler CNN that has also been used in numerous studies as a model of the ventral visual system (*65–68*). Together, our findings indicate that emotion selectivity can spontaneously emerge in CNN models trained to recognize visual objects, and these emotion-selective neurons play a significant role in recognizing emotion in natural images, lending credence to the notion that the visual system’s ability to represent affective information may be intrinsic.

### Affective processing in the visual cortex

The perception of opportunities and threats in complex visual scenes represents one of the main functions of the human visual system. The underlying neurophysiology is often studied by having observers view pictures varying in affective content. (*69*) reported greater functional activity in the visual cortex when subjects viewed pleasant and unpleasant pictures than neutral images. (*70*) showed the visual cortex has differential sensitivities in response to emotional stimuli compared to the amygdala. (*71*) demonstrated that emotional significance (e.g., valence or arousal) could modulate the perceptual encoding in the visual cortex. Two competing but not mutually exclusive groups of hypotheses have been advanced to account for emotion-specific modulations of activity in the visual cortex. The so-called reentry hypothesis states that the increased visual activation evoked by affective pictures results from reentrant feedback, meaning that signals arising in subcortical emotion processing structures such as the amygdala propagate to the visual cortex to facilitate the processing of motivationally salient stimuli (*72–74*). Recent work (*20*) provides support for this view. Using multivariate pattern analysis and functional connectivity, these authors showed that (1) different emotion categories (e.g., pleasant versus neutral and unpleasant versus neutral) are decodable based on the multivoxel patterns in the visual cortex and (2) the decoding accuracy is positively associated with the strength of connectivity from anterior emotion-modulating regions to ventral visual cortex. A second group of hypotheses states that the visual cortex may itself have the ability to code for the emotional qualities of a stimulus, without the necessity for recurrent processing (see (*75*) for a review). Evidence supporting this hypothesis comes from empirical studies in experimental animals (*76*, *77*) as well as in human observers (*78*), in which the extensive pairing of simple sensory cues such as tilted lines or sinusoidal gratings with emotionally relevant outcomes shapes early sensory responses (*79*). Beyond simple visual cues, recent computational work using deep neural networks has also suggested that the visual cortex may intrinsically represent emotional value as contained in complex visual media such as video clips of varying affective content (32). Our results, by showing that emotion-selective neurons exist in all layers of two CNN models of the visual cortex and that these neurons play an important role in emotion recognition, add to computational evidence that the visual cortex may have the intrinsic ability to assess the emotional significance of visual stimuli.

### Neural selectivity in ANNs and the brain

That CNNs, or more generally ANNs, can be trained to recognize a large variety of visual objects has long been recognized. Remarkably, recent studies note that ANNs trained on recognizing visual objects can spontaneously develop selectivity for other types of input, including visual numbers and faces (*80*). The number sense is considered an inherent ability of the brain to estimate the quantity of certain items in a visual set (*81*, *82*). There is significant evidence demonstrating that the number sense exists in both humans (e.g., adults and infants) (*83–85*) and non-human primates (e.g., numerically naïve monkeys) (*86–88*). (*89*) found that number-selective units spontaneously emerged in a deep artificial neural network trained on ImageNet for object recognition. (*90*) demonstrated that number selectivity can even arise spontaneously in randomly initialized deep neural networks without any training. Both studies focused on the last convolutional layers, in which the number-selective units were found, and they also demonstrated that the emergence of number-selective units could result from the weighted sum of both decreasing and increasing the activity of some units. In addition, it is well known that face-selective neurons exist in humans (*91*) and non-human primates. (*80*) showed that neurons in a randomly initialized deep neural network without training could selectively respond to faces, and the neurons in the deeper layers are more selective. (*92*) demonstrated that brain-like functional segregation can emerge spontaneously in deep neural networks trained on object recognition and face perception and proposed that the development of functional segregation of face recognition in the brain is a result of computational optimization in the cortex. Augmenting this rapidly growing literature, our study demonstrates that emotion selectivity can emerge in deep artificial neural network models of the human visual system trained to recognize objects.

### Layer dependence

Like the biological brain, the CNN model has a layered structure which allows the processing of information in a hierarchical fashion. Our layer-wise analysis showed that the extent and strength of emotion selectivity are a function of the model layers. Compared to the early layers, the deeper layers have larger portions of neurons that show emotion selectivity, and the selectivity is stronger, consistent with the previous observations that deeper layers of CNN models encode more abstract concepts. For example, (*40*, *93*) examined the internal representations of different layers in a CNN and found that deeper layers of the network tend to encode more abstract concepts, such as object parts and textures. The layered processing of emotional information may have several functional benefits. First, by processing visual information in hierarchical stages, the brain can quickly and efficiently respond to stimuli without the need for a complete and detailed analysis of the entire stimulus at once (*94–96*). This is especially important for the processing of emotionally salient stimuli, as quick and accurate emotional responses can be crucial for survival. Second, it would offer more flexibility for the processing of emotion at different levels of detail, which may depend on the perception task and the environmental context. For example, if the stimulus is perceived as significant or crucial for survival, it elicits a stronger and more widespread neural response, engaging multiple regions and processing stages. On the other hand, if the stimulus is not significant, it elicits a weaker and more limited neural response involving fewer regions or layers and processing stages (*97–99*). Third, the integration of information from different levels allows for a more complete and nuanced representation of the visual stimulus and emotional response. This allows for the creation of a final representation that takes into account not just the visual properties of the stimulus but also its emotional significance and its impact on the individual (*100–102*). Lastly, by processing information in a layer-dependent manner, the brain can adapt and change the processing of information based on experience and learning (*103*). This allows the brain to refine its processing strategies and improve its performance over time (*104*).

### Relation to prior literature

(*32*), to the best of our knowledge, is the first to examine emotion processing in deep neural networks. Their model, which is a modified AlexNet called the EmoNet, was shown to have the ability to classify affective images into 20 different emotion categories. Importantly, using a 20-way linear decoder, they further showed that neural activities in different layers of the network especially the deeper layers can differentiate different emotions in the input images, suggesting the existence of emotion selectivity neurons in CNNs. Building on this work, our main contributions are threefold: (1) confirming and characterizing emotion selectivity at the single filter (neuron) level, (2) demonstrating the functional significance of emotion-selective neurons through the application of lesion and attention enhancement methods, and (3) replicating the findings across two CNN models (VGG-16 and AlexNet) and two affective image sets (IAPS and NAPS).

### Limitations and other considerations

Several limitations of our study should be noted. Firstly, emotion was divided into three broad categories: pleasant, unpleasant, and neutral. While this is in line with many neurophysiological studies in humans, future work should examine finer differentiations of emotion, e.g., joy, sadness, horror, disgust, and so on, and their neural representations in the brain. Secondly, there might be other factors (e.g., low-level features) that drive the emotion selectivity of neurons. Since we used grayscale images in this study, we can rule out color as a possible confounding low-level feature. Applying the GIST algorithm (*105*) to extract low-level features from images and the support vector machine (SVM) algorithm (*106*), we found that images from different emotion categories cannot be decoded from the low-level features; see Figure S9 of Supplementary Materials. The impact of an image’s object category and its emotion category on neural activation was examined by placing images in the IAPS and NAPS datasets into object categories based on the descriptions of the images (Figure S11A and S12A of Supplementary Materials) and applying Two-Way ANOVA tests to filter activations in the VGG-16 model. We found that the neurons responded more strongly to emotion categories than object categories and there were significant interactions between the two categories in deeper layers (Figure S11A and S12A of Supplementary Materials). We do note that, as the number of images in different object categories are relatively small in both affective datasets, this analysis should be viewed as preliminary. The influence of other factors such as the presence of faces and image animacy is more difficult to ascertain. Thirdly, although the present study is motivated by neuroscience questions, to what extent our results have a direct bearing on understanding brain function is unclear. Whereas previous work did compare activities in VGG-16 and other deep neural networks with neural recordings during object recognition (Tiago Marques, Martin Schrimpf, and James J. DiCarlo 2021; Ratan Murty et al. 2021; Uran et al. 2022; Zhuang et al. 2021), there is no study to date comparing activities in deep neural networks and neural recordings during emotion recognition. In this sense, this work’s neural relevance should be considered speculative.

## Materials and Methods

### Affective picture sets

Two sets of widely used affective images were used in this study. The IAPS library includes 1,182 images covering approximately 20 subclasses of emotions such as joy, surprise, entrancement, sadness, romance, disgust, and fear. The NAPS library has 1,356 images that can be divided into similar subclasses. For both libraries, each image has a normative valence rating, ranging from 1 to 9, indicating whether the image expresses unpleasant, neutral, or pleasant emotions; the distributions of the valence rating from the two datasets were given in Fig.S1-C (right). In this study, for simplicity and following the common practice in human imaging studies of emotion (*20*, *107–109*), we classified images into three main categories based on their valence scores: “pleasant,” “neutral,” and “unpleasant.” For images that fell near the boundary between categories, we used soft thresholds of 4.3±0.5 and 6.0±0.5 to determine their classification as either “unpleasant” or “neutral,” or “neutral” or “pleasant.” We also visually examined each image to confirm its category. Finally, any images that we could not confidently classify were marked as “unknown” and removed from the analysis. This process resulted in some differences in the number of images in each category from the original datasets. After this categorization, IAPS images were divided into 296 pleasant, 390 neutral, and 341 unpleasant images, and NAPS images into 352 pleasant, 477 neutral, and 281 unpleasant images (see Fig.S1-B). These images were transformed from the original color images to grayscale images prior to the commencement of the study reported here. The goal was to remove color as a possible low-level visual feature confounding the emotion selectivity analysis.

### The convolutional neural network model

VGG-16, a well-tested deep convolutional neural network for natural image recognition, was used in this study to evaluate emotion selectivity. It has 13 convolutional layers followed by three fully connected layers, with the last fully connected layer containing 1000 units for recognizing 1000 different types of visual objects. Each layer of VGG-16 contains a large number of filters/channels, the application of each of which results in a feature map consisting of a large number of units. For convenience, and to stress neurobiological relevance, these filters/channels were often referred to as artificial neurons or simply neurons in this paper. Each neuron is characterized by a ReLU activation function (see Fig.S1-A). Through this function, neurons within a given layer, upon receiving and processing the input from the previous layer, yield activation maps (i.e., feature maps) which become the input for the next layer. Previous studies have compared the activation patterns of the VGG-16 model with experimental recordings from both humans and non-human primates and found that early layers of the model behave similarly to early visual areas such as V1, whereas deeper layers of the model are more analogous to higher-order visual areas such as the object-selective lateral occipital areas (*22*, *110–112*).

In this study, VGG-16 was used in two ways. First, to examine whether emotional selectivity emerges in neurons trained to recognize objects, we took the VGG-16 model pre-trained on 1.2 million natural images from the ImageNet, presented affective pictures from the two aforementioned affective picture datasets to the model, and analyzed the activation profiles of neurons from each layer. The emotional selectivity of each neuron was determined from these activation profiles (see below). Second, to test the functionality of the emotion-selective neurons, we replaced the last layer of the VGG-16 with a two-unit fully connected layer and trained the connections to this two-unit layer to recognize two categories of emotion: pleasant versus non-pleasant, neutral versus non-neutral, or unpleasant versus non-unpleasant. The training of the last two-unit emotion recognition layer used cross-entropy as the objective function. It is worth noting that, aside from the last emotion-recognition layer, the other layers’ weights in the VGG-16 network remained the same as that trained on the ImageNet data; in other words, they were frozen.

The training data and the testing data for the final 2-unit emotion recognition layer of our model were separate for IAPS and NAPS to avoid overfitting. Specifically, for each emotion category, we partitioned the images from both datasets into training, validation, and testing subsets at a ratio of 50%:25%:25%. We used a learning rate of 1𝑒 − 3, trained for 10 epochs, and set the batch size to 128. Finally, we employed the F1-score to assess the performance of our model in emotion recognition.

### Emotion selectivity definition

We used two methods to evaluate the differential responses of a neuron to images from different emotion categories (pleasant, neutral, or unpleasant). Tuning value emphasizes the normalized response to images from the same category. It is used in Figure 2 to illustrate possible response profiles or tuning curves of different neurons. The selective index (SI), in contrast, emphasizes the difference between responses to images from one emotion category and those from other emotion categories. It is thus more suitable for quantifying the emotion selectivity of a neuron. Results reported in Figures 3-4 as well as in Supplementary Materials were done with the SI.

#### Tuning value calculation

We followed the method in (*43*) for calculating the tuning value in Figure 2. The tuning v focuses on the strength or magnitude of a neuron’s response to a particular emotion, relative to its average response. The details can be found below.

The output from each filter also referred to as a neuron in this study, see Fig.S1-A, can be written as:

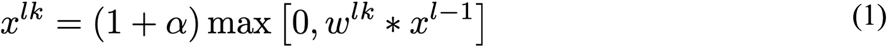

where *w^lk^* indicates the weights of the *k^th^* filter in the *l^th^* convolutional layer, and ∗ indicates mathematical convolution which applies matrix multiplication between *w* and the outputs *X* from the (*l* − 1)*^th^* layer. Of note in Eq. (1) is that the ReLU activation function typically has a slope of 1 (𝛼 = 0). Here in this work, the slope is a tunable parameter. By tuning the slope of the ReLU function, we change the gain of the neuron, simulating the effect of feature-based attention control (*43*, *53*).

Let 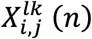 represents the response of the unit located at coordinates (𝑖, 𝑗) in the *k^th^* filter in layer *l* to image *n*. Then

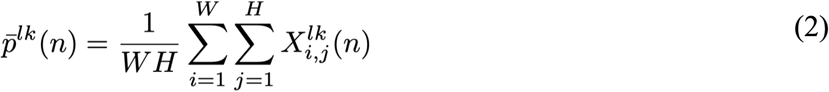

is the response to the image averaged across the entire filter. Here *W* and *H* represent the width and height of the feature map. Thus, the mean activity of the filter *k* in layer *l* in response to all images in a dataset can be formulated as:

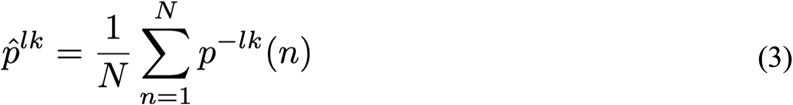

where *N* represents the total number of images in a given set. The tuning value of the filter is calculated according to

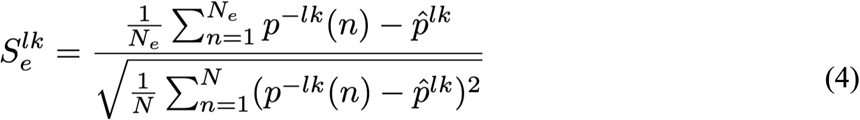

where 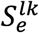 represents the normalized activation of filter *k* in layer *l* in response to all images of emotion category *e*, where *e* ∈ {*pleasant*, *neutral*, *unpleasant*}. A neuron is considered selective for a specific emotion if the normalized activation for the images within that emotion category is highest among the three possible values. For example, if 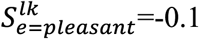, 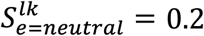, and 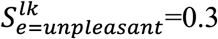, the artificial neuron *k* is considered selective for “unpleasant images”.

#### Selectivity index calculation

Selectivity Index (SI) (*113*) is defined as follows. First, consider

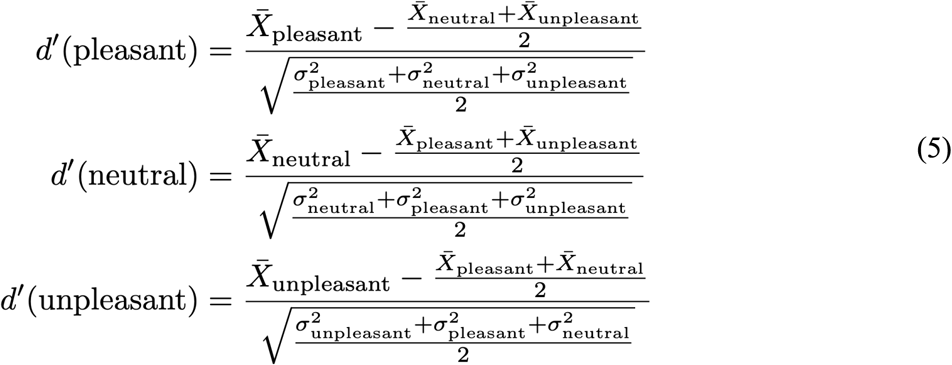

where X*_pleasant_*, X*_neutral_*, X*_unpleasant_* represents the mean response to the pleasant, neutral, and unpleasant categories, respectively; 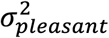, 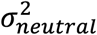, and 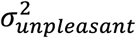 represents the variance of the response to the pleasant, neutral, and unpleasant category, respectively. SI is the largest *d*′ and the emotion that gives rise to the largest *d*′ defines the emotion for which the neuron is selective.

#### Identification of emotion-selective neurons

To guard against spurious identification of emotion selectivity and ensure that neurons designated to be selective for an emotion do so for both datasets, we applied two analyses. First, we rank-ordered neurons according to their SI values, eliminated neurons with small SI values, and tested the emotion recognition performance under attention enhancement of the remaining neurons (see below). Increasing the percentage of neurons eliminated until we saw a significant change in performance. That percentage was then defined as the threshold for defining emotion selectivity within a dataset (see Figure 3C for an example of finding the threshold for the pleasant category on the IAPS dataset). Second, for neurons identified as selective for certain emotions based on IAPS and that based on NAPS, we overlapped the two sets of neurons and considered the overlapped neurons to be the genuine emotion-selective neurons.

### Testing the functionality of the emotion-selective neurons

Do the emotion-selective neurons defined above have a functional role? We applied two different approaches to examine this question: lesion and attention enhancement.

#### Lesion

If the emotion-selective neurons are functionally important, then lesioning these neurons should lead to degraded performance in recognizing the emotion of a given image. Here the lesion of a specific neuron is achieved by setting its output to 0 (namely, setting 𝛼 = −1 in Eq. (1)). In our experiments, we lesioned the neurons selective for a given emotion as well as randomly selected neurons in a particular layer and observed the changes in the emotion recognition performance of the model.

#### Attention enhancement

We further tested whether enhancing the activity of an emotion-selective neuron can lead to performance improvement in emotion recognition. Following (*43*), the strength of α was increased from 0 to 5 with interval step size 0.1, where 𝛼 = 0 is the conventional choice and 𝛼 > 0 represents increased neuronal gain (i.e., enhanced feature attention). According to the feature similarity gain theory, increasing the gain of a neuron leads to enhanced performance of the neuron in perceiving stimuli with the relevant features. In our experiments, we enhanced the neurons selective for a given emotion as well as randomly selected neurons in a particular layer and observed the changes in the emotion recognition performance of the model (*43*) (see Fig.S2-A,B).

## Supporting information

**S1 Fig. Model development details**.

**S2 Fig. The activation of calculation with convolution and *ReLU***.

**S3 Fig. Selective Index.**

**S4 Fig. Number of selective neurons across layers by emotion category in dataset IAPS (Top) and NAPS (Bottom).**

**S5 Fig. Functional generalization analysis.**

**S6 Fig. Selective Index Quality(A) and Generalizability in AlexNet (B) of Emotion Selectivity across Two Datasets.**

**S7 Fig. Effects of enhancing emotion-selective neurons and randomly selected neurons in AlexNet.**

**S8 Fig. The functionality of selective neurons in AlexNet when performing lesion their responses to three categorical emotions: pleasant, neutral, and unpleasant**

**Table S1 Parallel Results Produced with VGG-16 and AlexNet.**

**Table S2 Original and Enhanced Performance (F1-score) in VGG-16 and AlexNet.**

**S9 Fig. Pairwise decoding results using low-level features (GIST).**

**S10 Fig. Number of images involving faces in top 100 images that evoked the strongest response of selective neurons.**

**Table S3 Valence and Arousal of Top 100 Images Across Selective Neuron Categories.**

## Author contributions

**Conceptualization:** Peng Liu, Mingzhou Ding, Ruogu Fang

**Data Curation:** Peng Liu, Ke Bo

**Formal Analysis:** Peng Liu

**Funding Acquisition:** Peng Liu, Mingzhou Ding, Ruogu Fang

**Investigation:** Peng Liu, Ke Bo, Mingzhou Ding, Ruogu Fang

**Methodology:** Peng Liu, Ke Bo, Mingzhou Ding, Ruogu Fang

**Project Administration:** Mingzhou Ding, Ruogu Fang

**Resources:** Mingzhou Ding, Ruogu Fang

**Software:** Peng Liu

**Supervision:** Mingzhou Ding, Ruogu Fang

**Validation:** Peng Liu, Ke Bo, Mingzhou Ding, Ruogu Fang

**Visualization:** Peng Liu, Ke Bo

**Writing – Original Draft Preparation:** Peng Liu, Mingzhou Ding, Ruogu Fang

**Writing – Review & Editing:** Peng Liu, Ke Bo, Mingzhou Ding, Ruogu Fang

**Competing interests:** The authors declare that they have no competing interests.

**Data and materials availability:** All data are publicly available in the main text or the supplementary materials. The analysis code will be available and may be requested from the authors.

## Supplementary Materials

Eight topics related to the study reported in the main manuscript are addressed in this Supplementary Materials.

### Topic 1. Additional details of model developments, image datasets and methods

The structure of artificial neurons, the number of images in each dataset, and valence distribution in each dataset are shown in Figure S1. The information flow of applying convolution and ReLU function on an input, the details of how to enhance and lesion artificial neurons, and the datasets and networks used in the study are illustrated in Figure S2.

**Figure S1.**
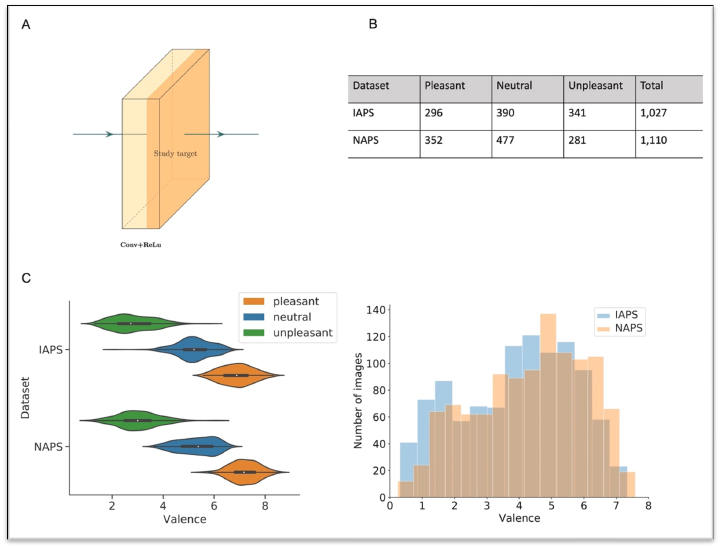
Model development details and image datasets. (**A**) Each convolutional layer is followed by one *ReLU* layer, the output of which reflects the responses of the artificial neurons the convolutional layer. Thus, in this study, the output of the *ReLU* layer is our study target for understanding the activity of the artificial neurons. (**B**) It shows the number of images of each emotion category in the two datasets used in this study. Two datasets were treated equally for defining emotion-selective neurons and related lesion and attention manipulations. (**C**) It shows how the divided categorial images match the valence score originally rated by human subjects in the two datasets. The C (left) shows the valence score distribution and the boundary score between the pleasant and neutral category: 4.3±0.5 and between the neutral and unpleasant category: 6.0 ±0.5. The C (right) shows the number of images per valence score across two datasets. Basically, this figure illustrates the details of the model development and the affective image datasets.

**Figure S2.**
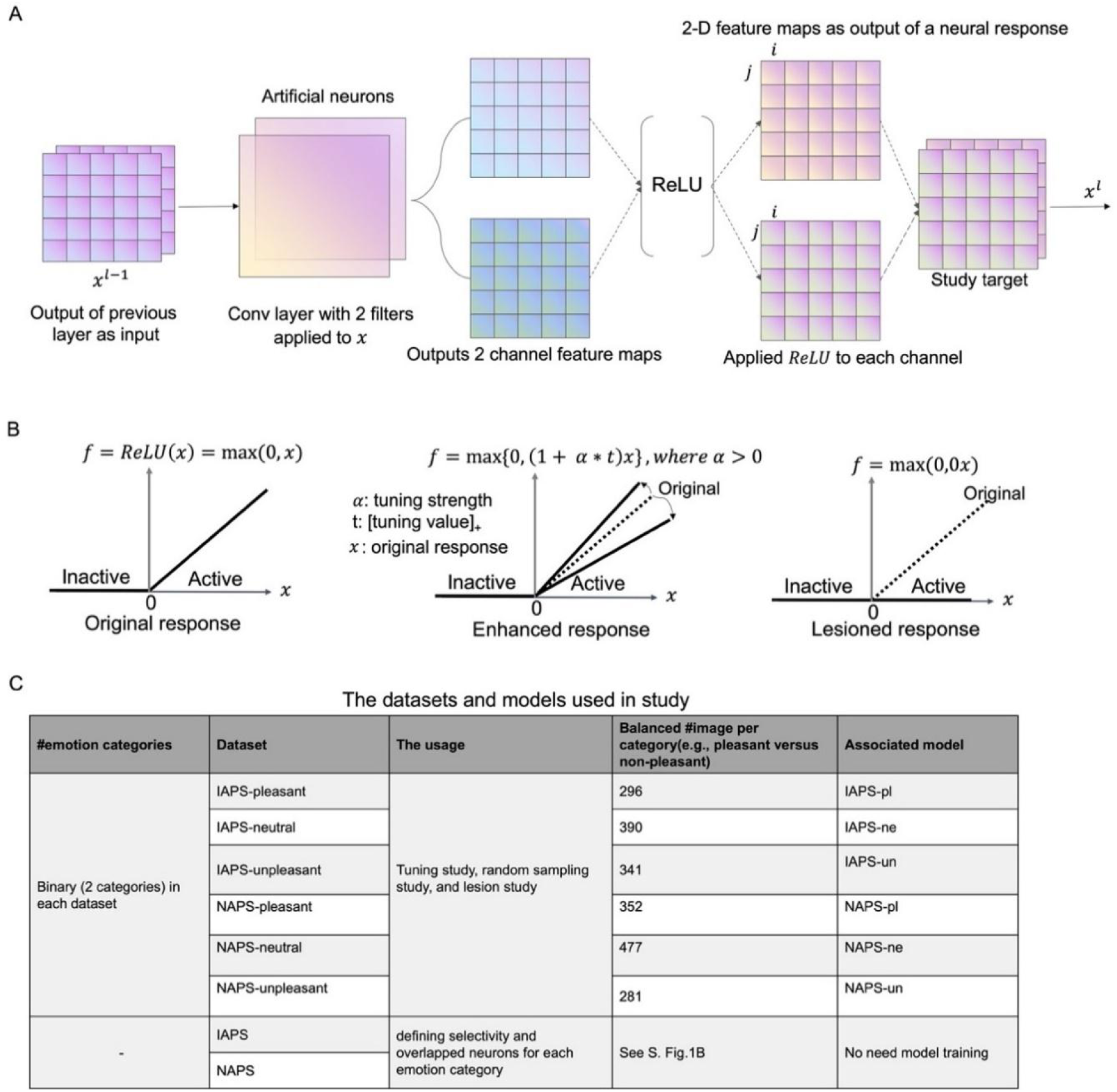
Further methodological details. (**A**) It shows that the information flow of applying convolution and *ReLU* operation to the input 𝑥 in layer 𝑙, which represents all the feature maps from the previous layer 𝑙 − 1. In this illustration, 𝑥 is composed of two channels of feature maps. Two filters (referred to as artificial neurons) in the following convolutional layer are applied to 𝑥 separately. Each filter convolution results in a channel of a new feature map, which then is passed through an activation function, *ReLU* = max(0, 𝑥). *ReLU* function indicates which features after the convolutional operation is activated or inactivated. The active units are valued with positive numbers, and the inactive units are valued with zeros by the *ReLU*. Fundamentally, this figure indicates how an artificial neuron responds to a stimulus and how the response activations are calculated in a CNN. (**B**) The bold black line represents the *ReLU* response (activation) values, and dash lines in the middle and right sub-figures represent the normal activation value of the *ReLU*. Three cases of activation behavior were investigated empirically. The normal excitation (left) is applying the original *ReLU*; the attention enhanced excitation (middle) is applying a positive weight 𝛼 to the activation value 𝑥; the inhabited activation through lesion is setting the activation values to be zeros instead, which performs like a lesion study. (**C**) It summarizes the datasets and the models used in the study. The number of images in the binary model is balanced. The non-* category images were randomly selected from another two categories.

### Topic 2. Additional analysis of the selective index

We examined the distribution and correlation of the selective index (SI) of selective neurons defined on dataset IAPS and NAPS, separately, in each layer and the number of selective neurons across layers by emotion category in IAPS and NAPS. The purpose is to answer the following questions: how many selective neurons there are in the network, how they depend on layers, and how the strength of selectivity depends on layers. The overall selective index is around 0.2 for IAPS and 0.15 for NAPS shown in Figure S3. The correlation between IAPS-defined emotion SI and NAPS-defined SI was computed and the result was shown in Figure S4. The left scatter plot, where neurons from all layers are combined, indicates a positive correlation (Pearson coefficient of 0.30) between IAPS- and NAPS-defined SI. The right plot shows that the correlation between the two SI indices increases as we move deeper into the network. This analysis further supports our claim that emotion selectivity is generalizable across the two datasets.

**Figure S3.**
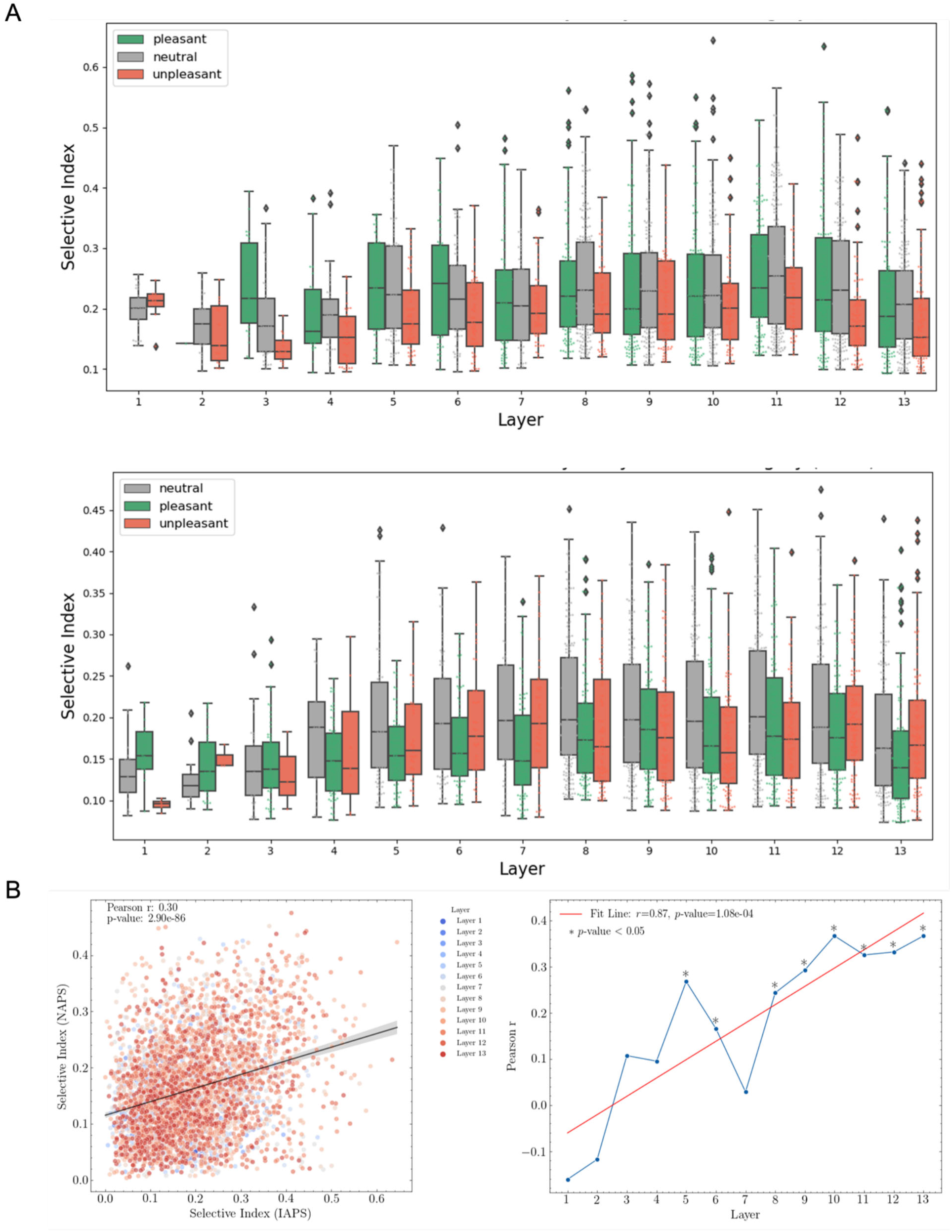
Additional analysis of selective Index. **(A)** Distribution of selective indices across layers by emotion category in dataset IAPS (Top) and NAPS (Bottom). **(B)** The correlation between IAPS-defined SI and NAPS-defined SI. (Left) Neurons from all layers are combined. (Right) The layer-wise correlation was plotted.

**Figure S4.**
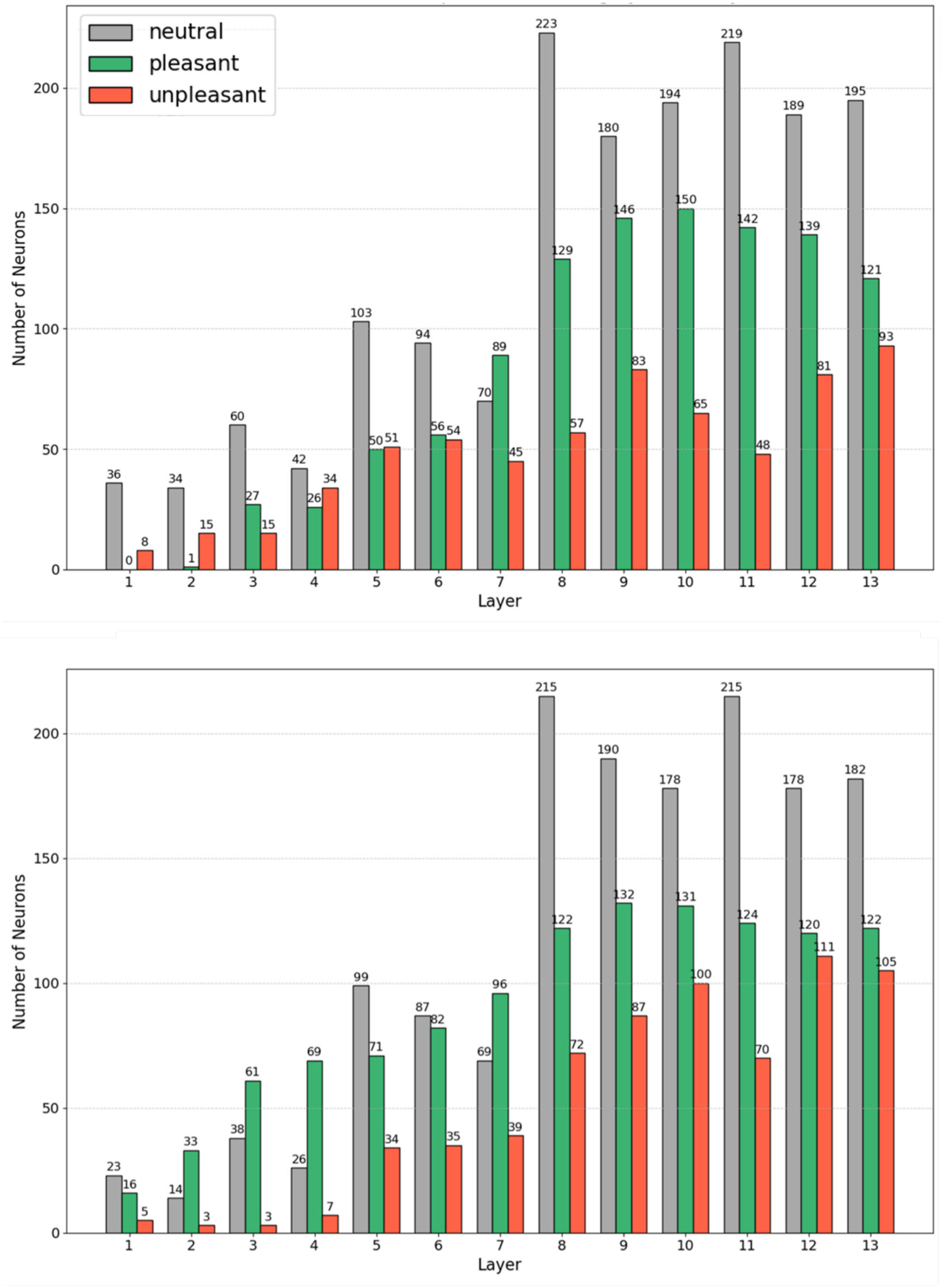
Number of selective neurons across layers by emotion category in dataset IAPS (Top) and NAPS (Bottom).

### Topic 3. Generalizability of emotion selectivity

We examined the functional generalization of selective neurons defined on IAPS and NAPS, separately, in Figure S5. The purpose is to further verify whether the emotion selectivity defined on one dataset can be functionally generalized to another dataset. The result is consistent with other results obtained by enchaining on selective neurons with their selective index defined on the same dataset (either IAPS or NPAS). It further supports our claim that emotion selectivity shares a functional property between the two datasets.

**Figure S5.**
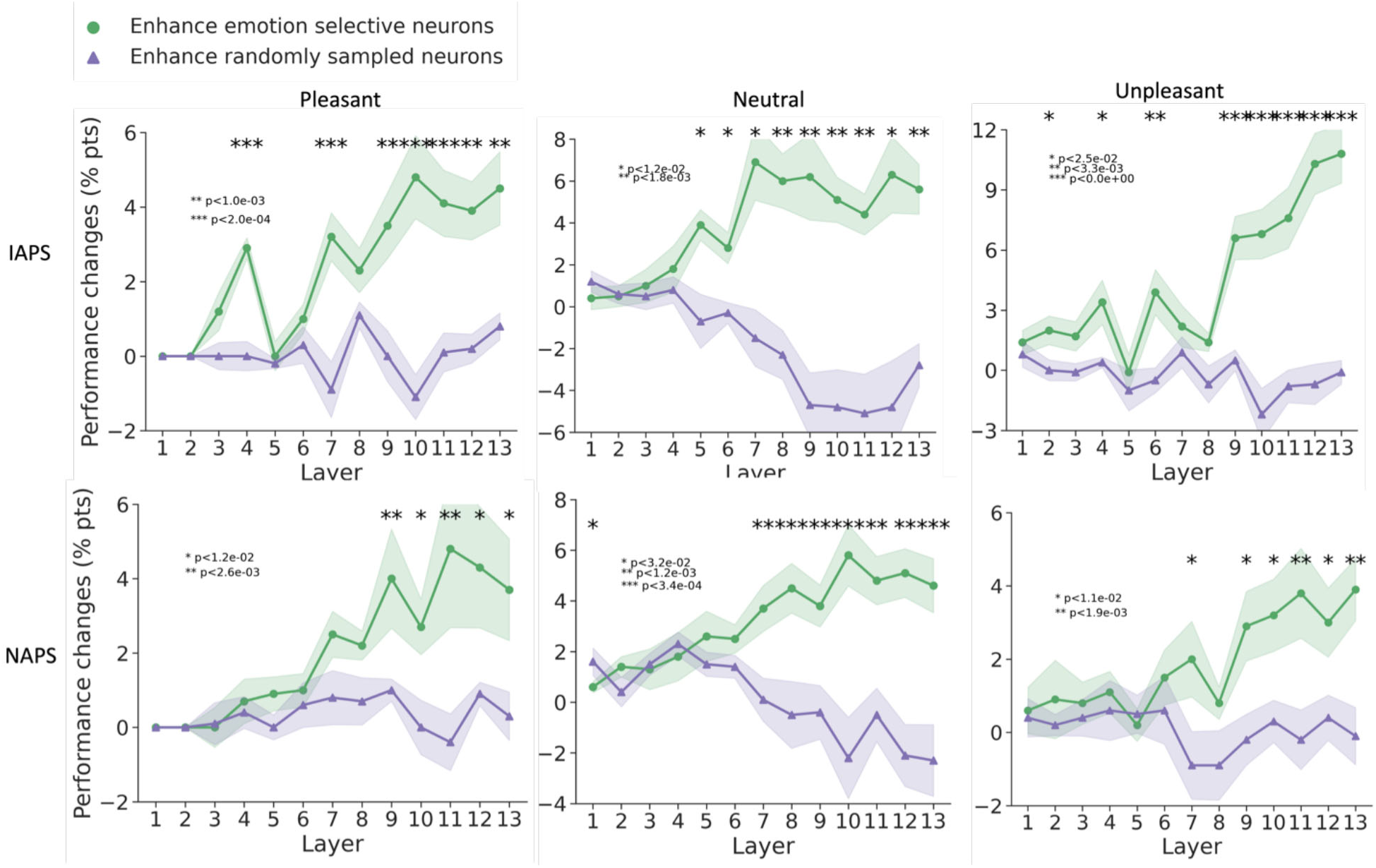
Functional generalization analysis. (**Top**) We analyzed the enhancement of emotion-selective neurons (defined post-threshold) versus random neurons in each VGG-16 layer trained on IAPS. The selective index was derived from NAPS and tested on IAPS. (**Bottom**) We analyzed the enhancement of emotion-selective neurons (defined post-threshold) versus random neurons in each VGG-16 layer trained on NAPS. The selective index was derived from IAPS and tested on NAPS.

### Topic 4. Result replication in AlexNet

We replicated the results in another network, AlexNet, shown in Figure S6, Figure S7, and Figure S8. The purpose is to demonstrate that the emergence of emotion selectivity is not an idiosyncratic property of a specific deep neural network. We summarized the parallel results produced with VGG-16 and AlexNet in Table S1 and Table S2.

**Figure S6.**
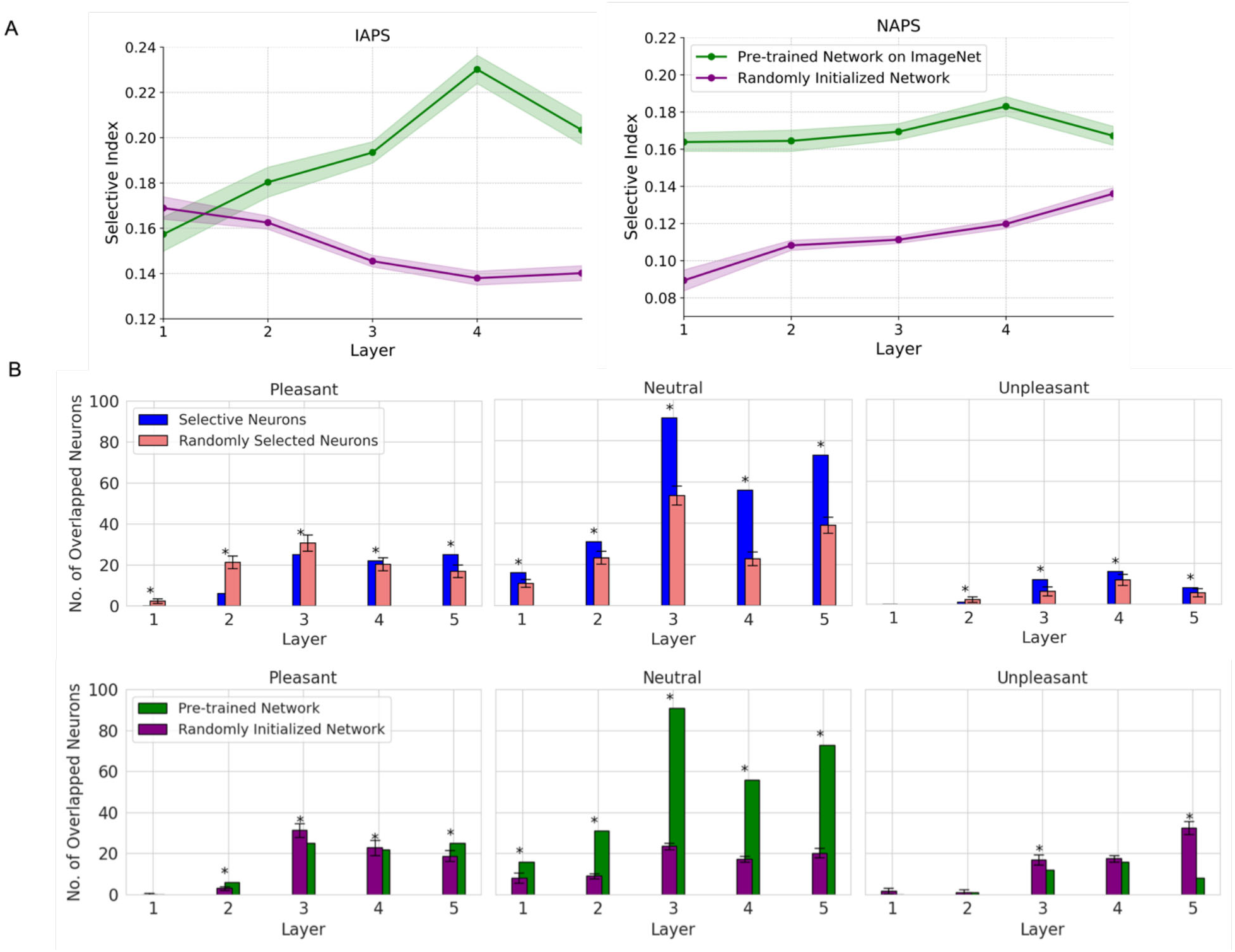
Selective Index Quality (A) and Generalizability in AlexNet (B) of emotion selectivity across two datasets. The comparison of number of overlapped neurons derived from selective neurons and randomly selected neurons is plotted (B-top). The one of number of overlapped neurons derived from pre-trained AlexNet on ImageNet and initialized AlexNet network with random weights (B-bottom). The goal of this comparison is to demonstrate the significance of learned features from ImageNet in developing neuron selectivity. However, merely counting the overlapping neurons might not be adequate; we should also take into account the selectivity index quality. This is particularly important when the total number of neurons in a layer is small.

**Figure S7.**
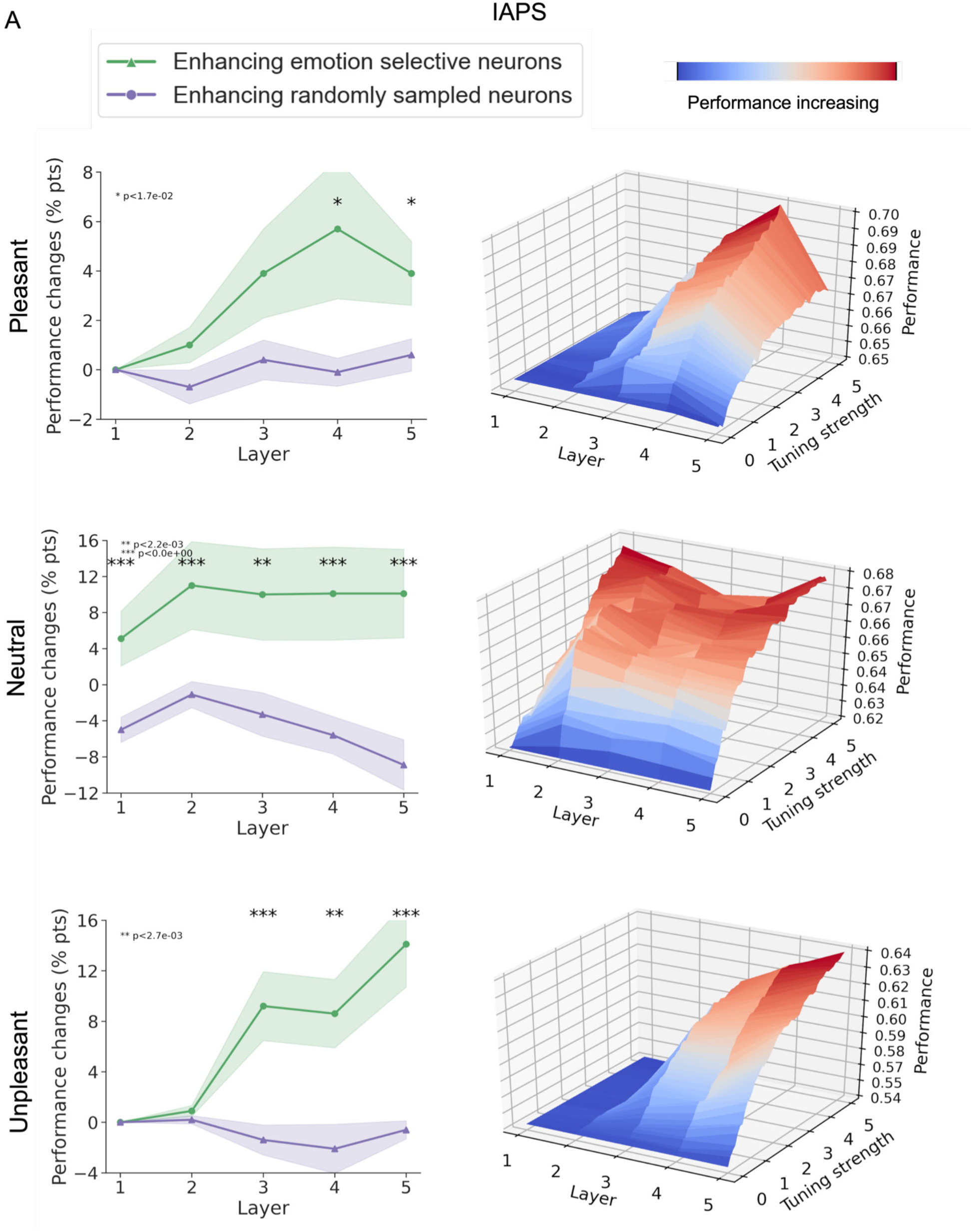
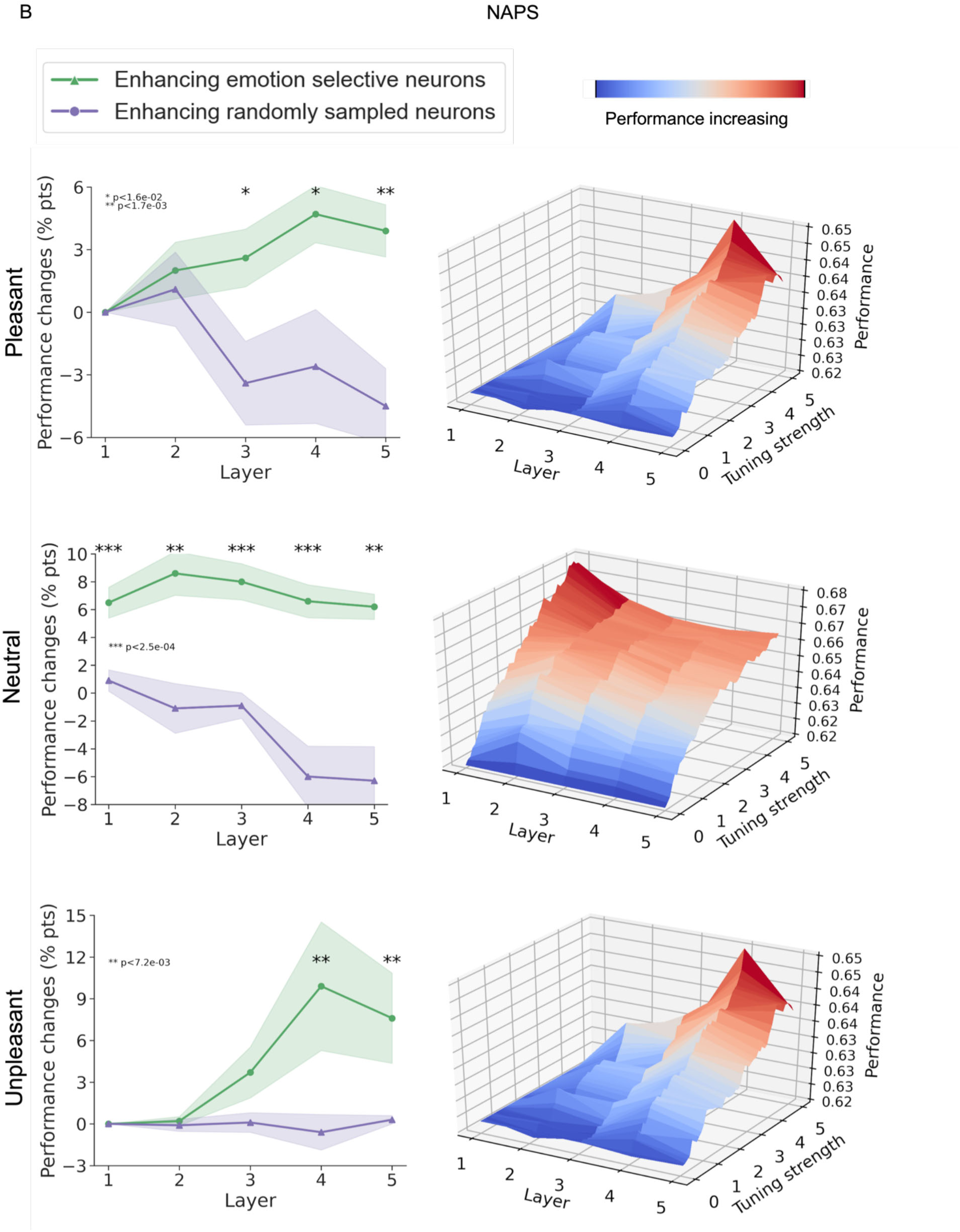
Effects of attention-enhancing emotion-selective neurons and randomly selected neurons in AlexNet. (**A**) IAPS dataset. (**B**) NAPS dataset.

**Figure S8.**
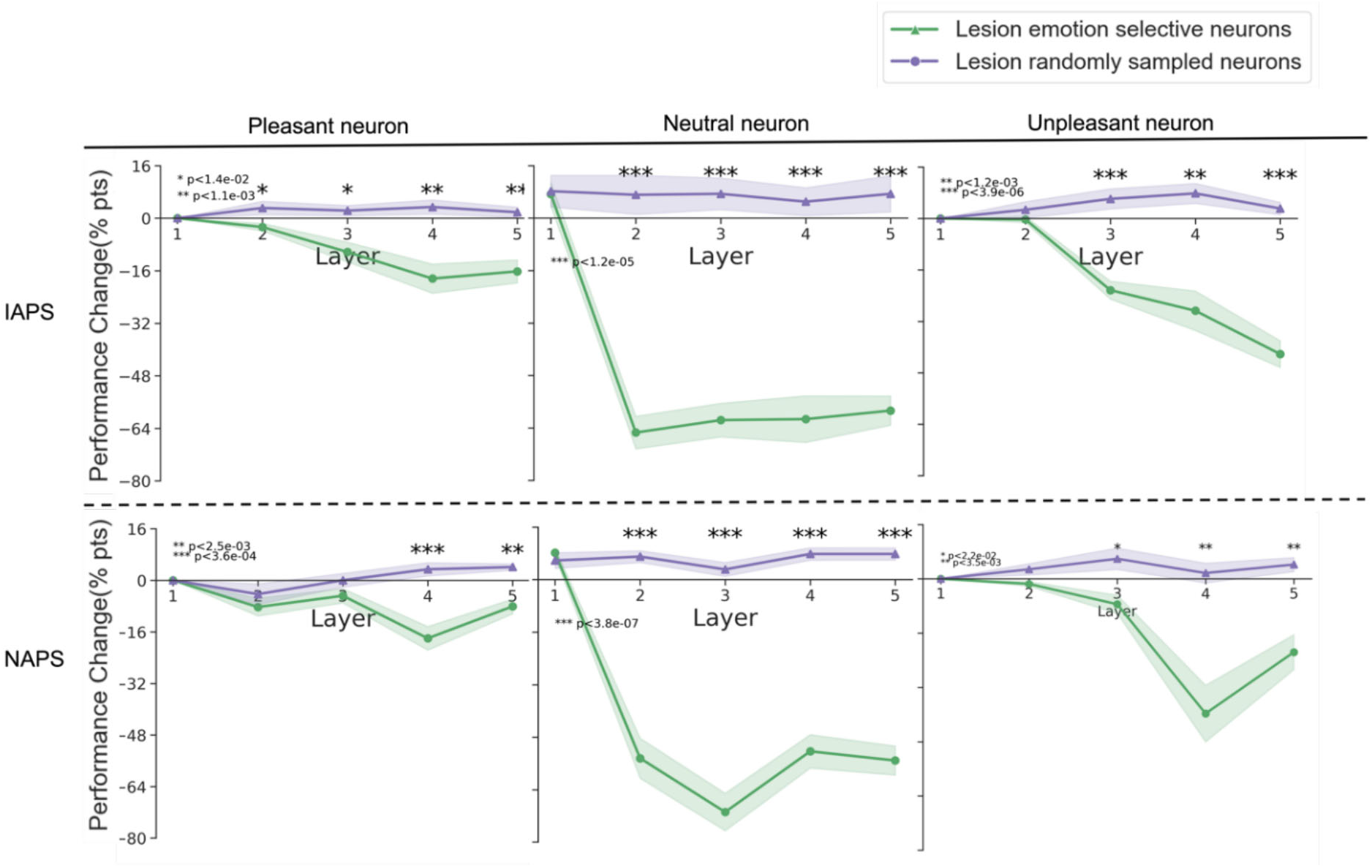
Effects of lesioning emotion-selective neurons and randomly selected neurons in AlexNet. (**A**) IAPS dataset. (**B**) NAPS dataset.

**Table S1.**
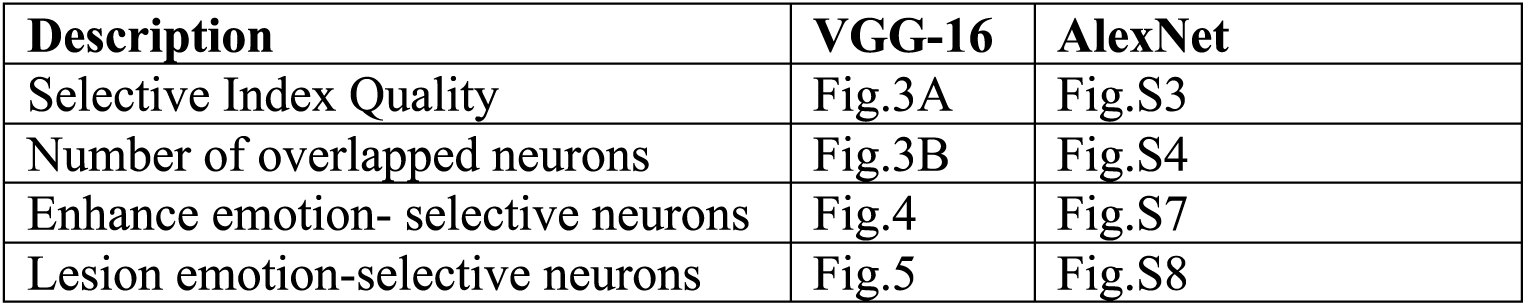
Results comparison between VGG-16 and AlexNet.

**Table S2.**
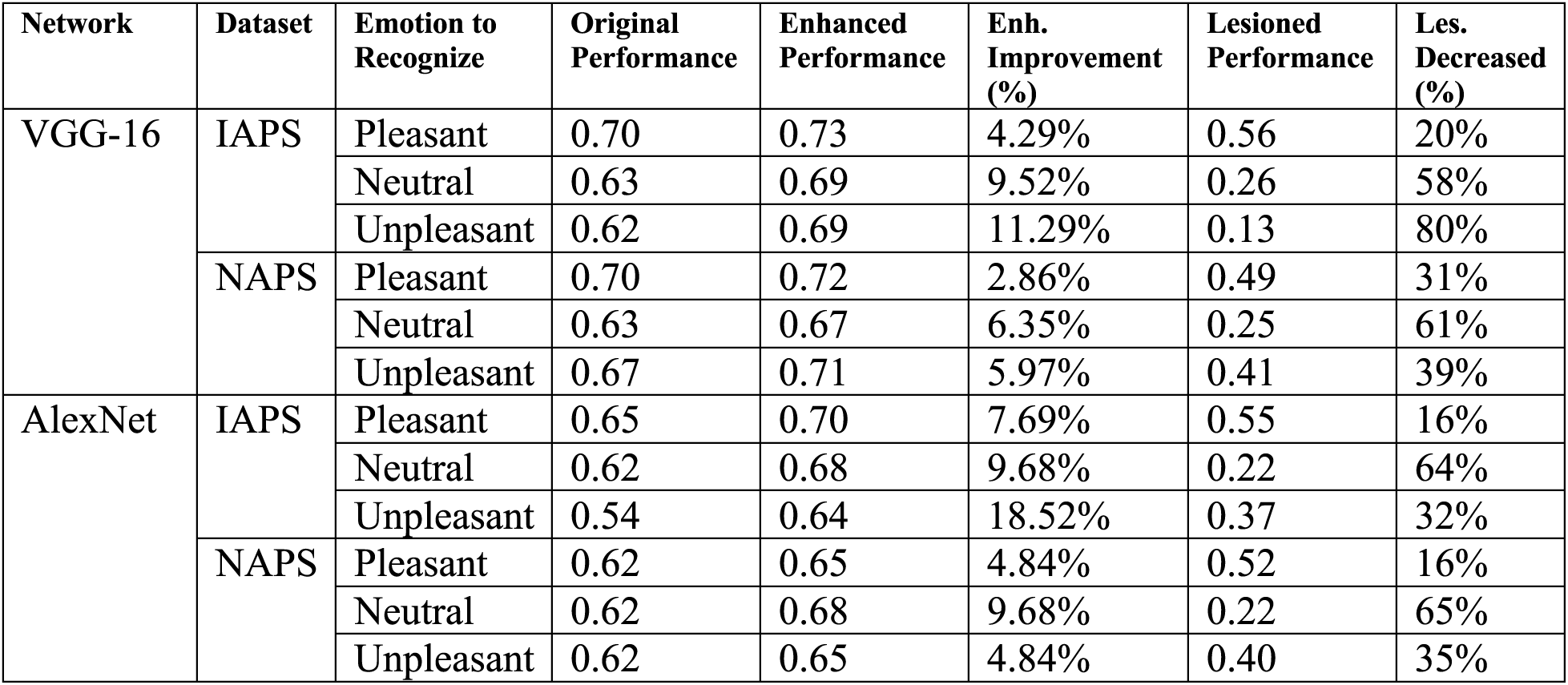
Original and Enhanced Performance (F1-score) in VGG-16 and AlexNet.

### Topic 5. Low-level features as possible confounding factors

low-level features were extracted from the images by using GIST algorithm. Pairwise emotion decoding was performed using (see Figure S9) using SVM. The mean accuracy for both IAPS and NAPS datasets approximates the chance level, suggesting that low-level GIST features are insufficient for decoding emotion categories from images.

**Figure S9.**
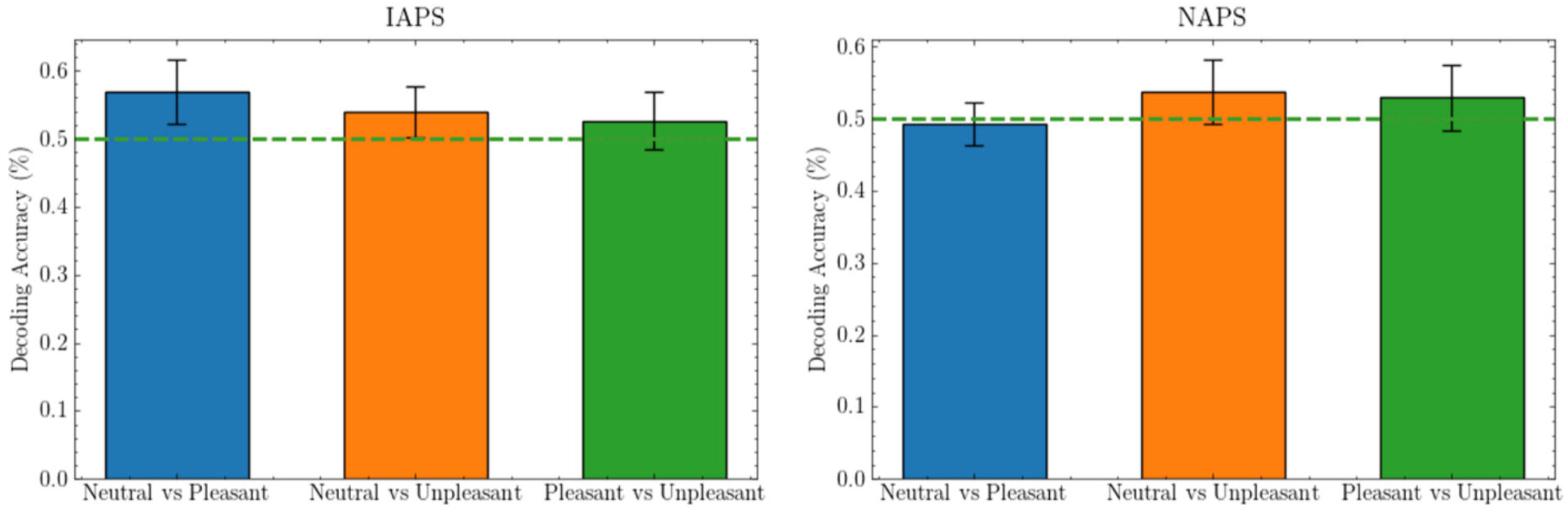
Pairwise decoding results using low-level features (GIST). Dash line indicates the chance level performance (50%). The dashed lines indicate chance-level performance and bars representing the average accuracy across 10 iterations of 5-fold cross-validation.

### Topic 6. Faces as possible confounding factors

The percentages of images involving faces in top 100 images (ranking based on neurons’ activation to each image) that evoked the strongest response of selective neurons for each emotion category (see Figure S10) are: 16%, 62%, and 30% for pleasant, neutral, and unpleasant, separately. The analysis demonstrates that development of emotion selectivity in these neurons is unlikely affected by potential facial encoding that might arise during the training of the network on ImageNet.

**Figure S10.**
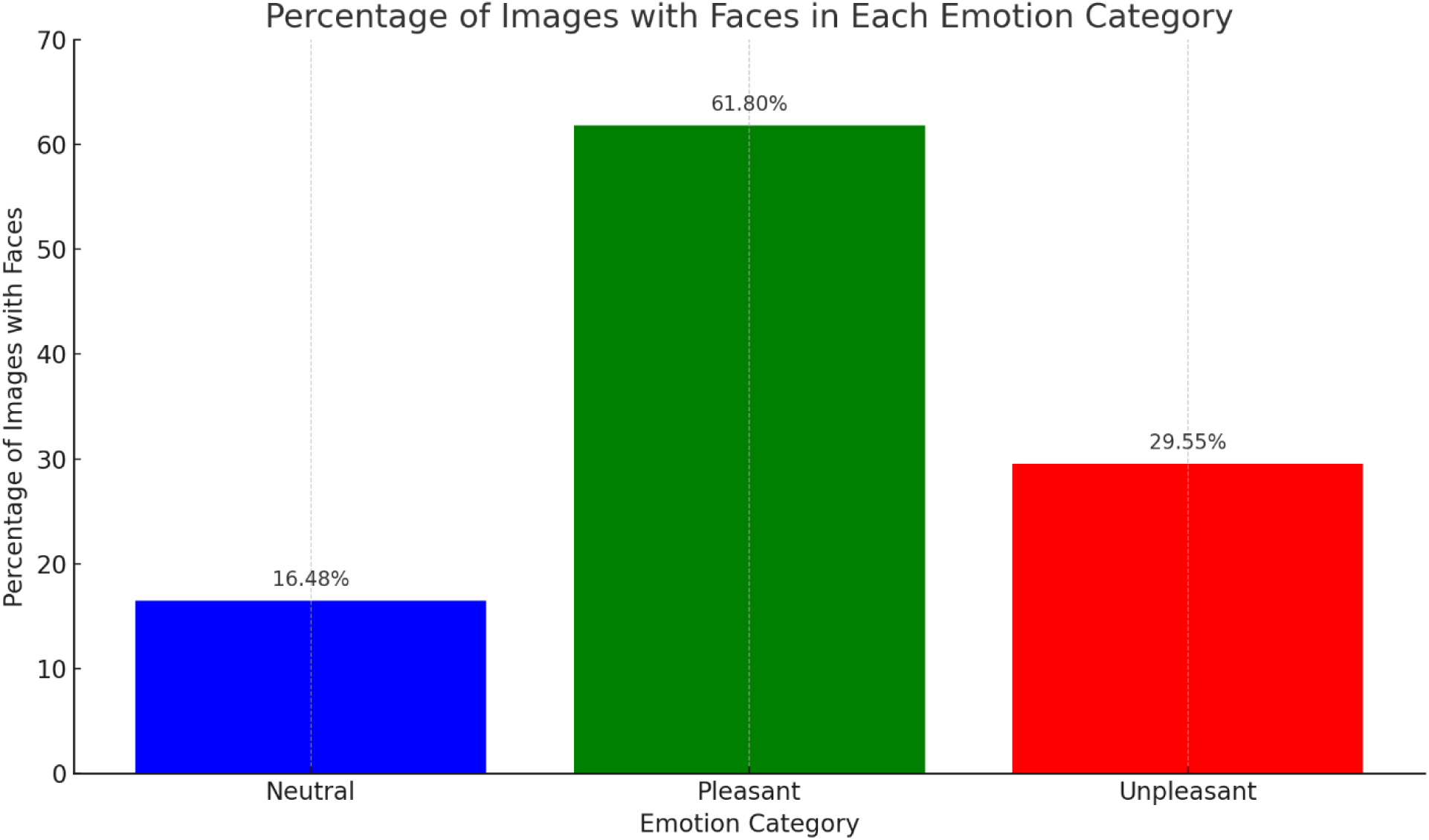
Number of images involving faces in top 100 images that evoked the strongest response of emotion-selective neurons.

### Topic 7. Animacy as possible confounding factors

#Table S3 shows the mean valence and arousal of the top 100 images across selective neuron categories. The purpose of this analysis is to estimate how much valence and arousal relevant to the images evoked by the selective neurons are captured. The result shows the mean valence: 6.770, 5.180, and 2.898 and mean arousal: 5.055, 3.970, and 5.816 for top images that evoked the strongest responses in neurons selective for pleasant, neutral, and unpleasant emotion. More importantly, these images appear to contain both animate and inanimate content, suggesting that animacy might not be a confounding factor.

**Table S3.**
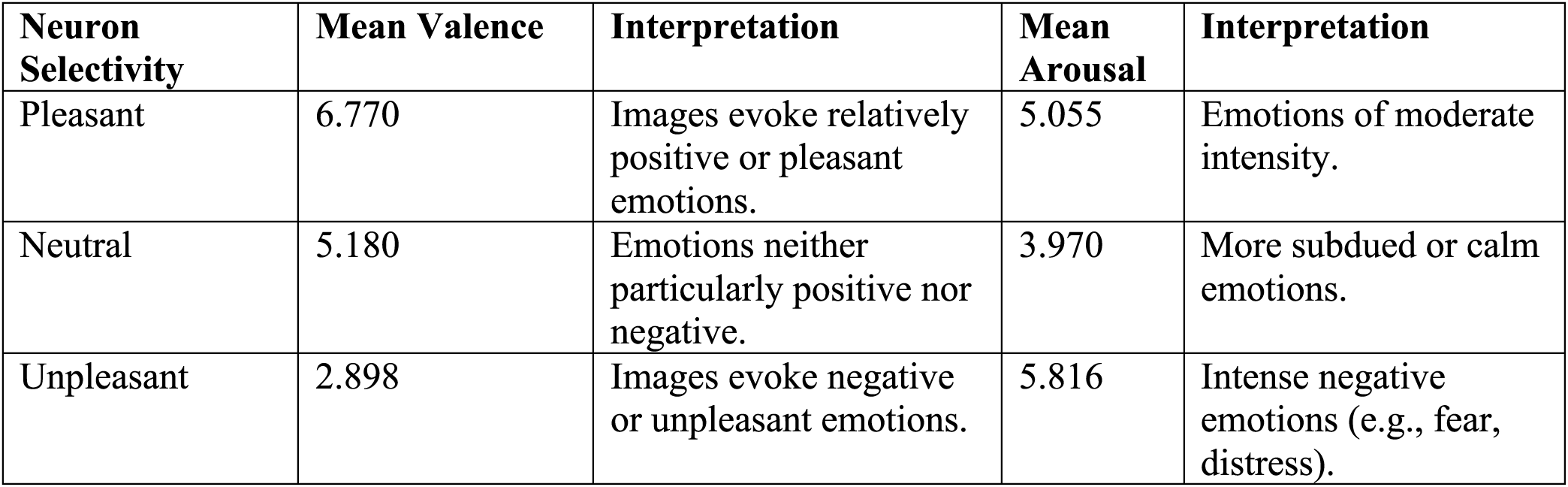
Valence and Arousal of Top 100 Images Across Selective Neuron Categories. Note: Image ranking is based on neurons’ activation to each image.

### Topic 8. Effects of emotion, object category, and their interaction on neuronal responses

As illustrated in Figures S11-S12, we examined, using a Two-Way ANOVA analyses, the impact of image emotion, image category, and their interaction on neuronal response. Object categories were identified based on the descriptions in the original datasets (refer to Figure S11A and S12A). Our findings reveal that the emotion category markedly affects neuronal activity in layers subsequent to the fifth (refer to Figure S11B-top and S12B-top), and the influence of the object category is increasing with layer depth but not significant. The interaction is significant in some deeper layers (refer to Figure S11B-bottom and S12B-bottom). It should be noted that this analysis should be viewed as preliminary, because the number of images in each object category is rather small, which may impact the analysis adversely. In addition, the selection of the images is also dataset-specific. For example, in the IAPS dataset, 15 images of dogs predominantly express negative emotions, whereas in the NAPS dataset, 35 images of dogs represent a mix of negative and positive emotions (see Figure S13). This variance indicates the necessity for additional studies to comprehensively understand the interaction between image emotion and category.

**Figure S11.**
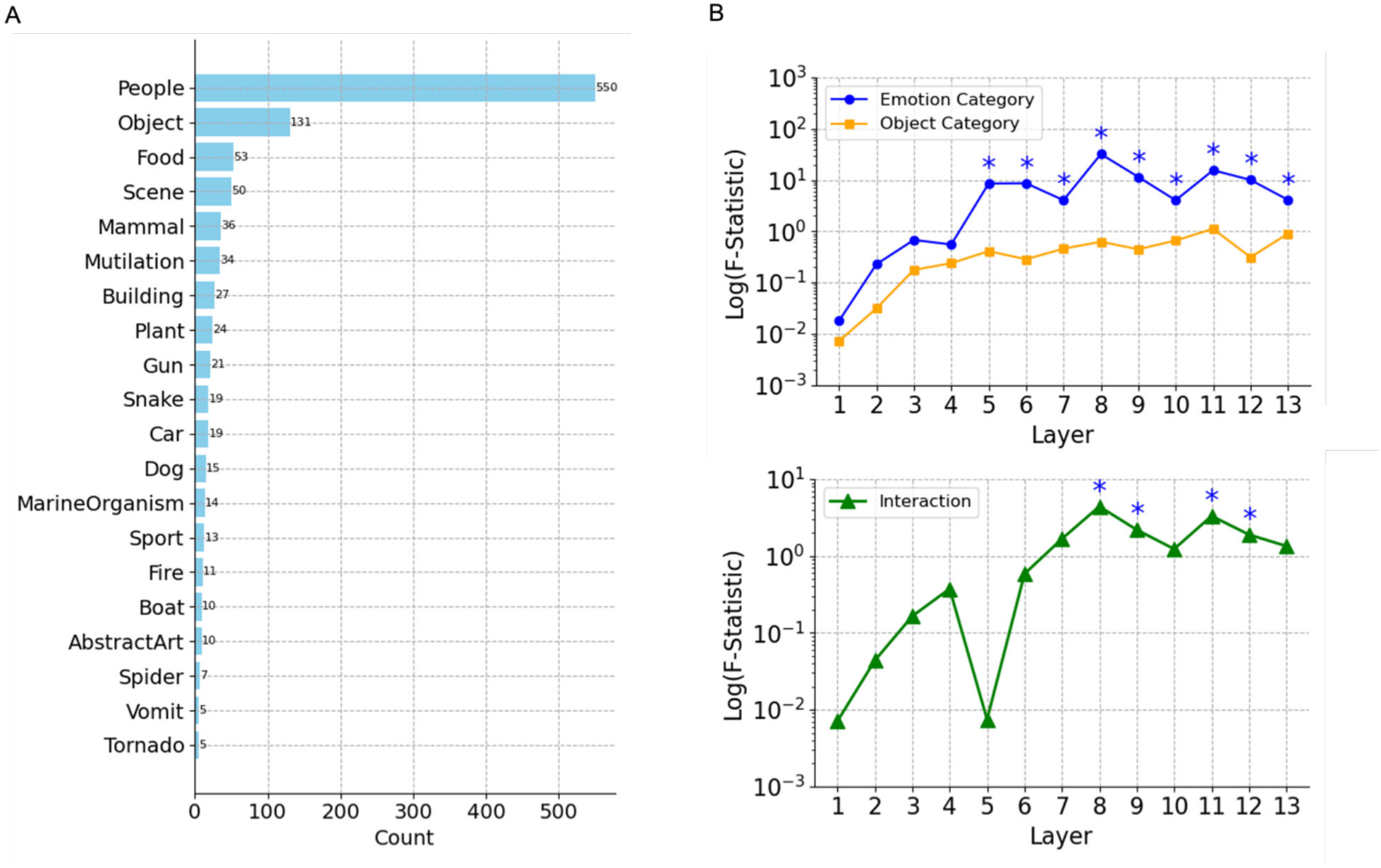
Effects of emotion and object category on filter activity using *IAPS* images. **A.** The number of images in each of the top 20 object categories. **B. (top)** The F-statistic (log scale) of the effect of emotion and object category on filter activations across layers of the VGG-16 neural network. The statistics are obtained from a Two-Way ANOVA test, where the dependent variable is the filter activity in response to images. The plot reveals how each factor impacts the filter responses and how this influence changes from the input to deeper layers of the network; (**bottom**) The F-statistic (log scale) of the interaction between emotion dependent filter activation and object category dependent filter activation. The statistics are obtained from a Two-Way ANOVA test. *\* indicates the influence is statistically significant*.

**Figure S12.**
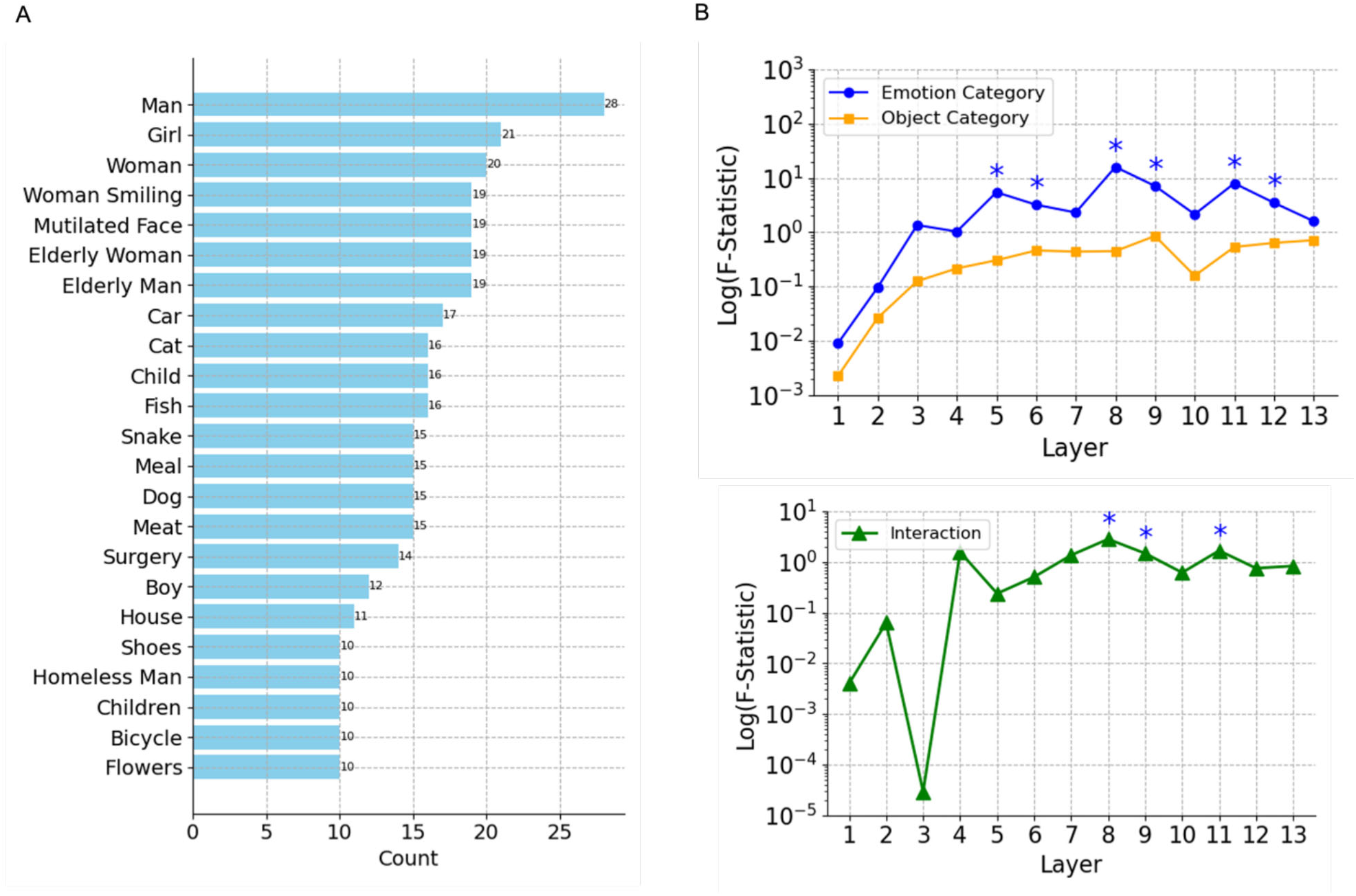
Effects of emotion and object category on filter activity using *NAPS* images. **A.** The number of images in each of the top 23 object categories. **B. (top)** The F-statistic (log scale) of the effect of emotion and object category in filter activations across layers of the VGG-16 neural network. The statistics are obtained from a Two-Way ANOVA test, where the dependent variable is the filter activity in response to images. The plot reveals how each factor impacts the filter responses and how this influence changes from the input to deeper layers of the network; (**bottom**) The F-statistic (log scale) of interaction between emotion and object category. The statistics are obtained from a Two-Way ANOVA test. *\* indicates the influence is statistically significant*.

**Figure S13.**
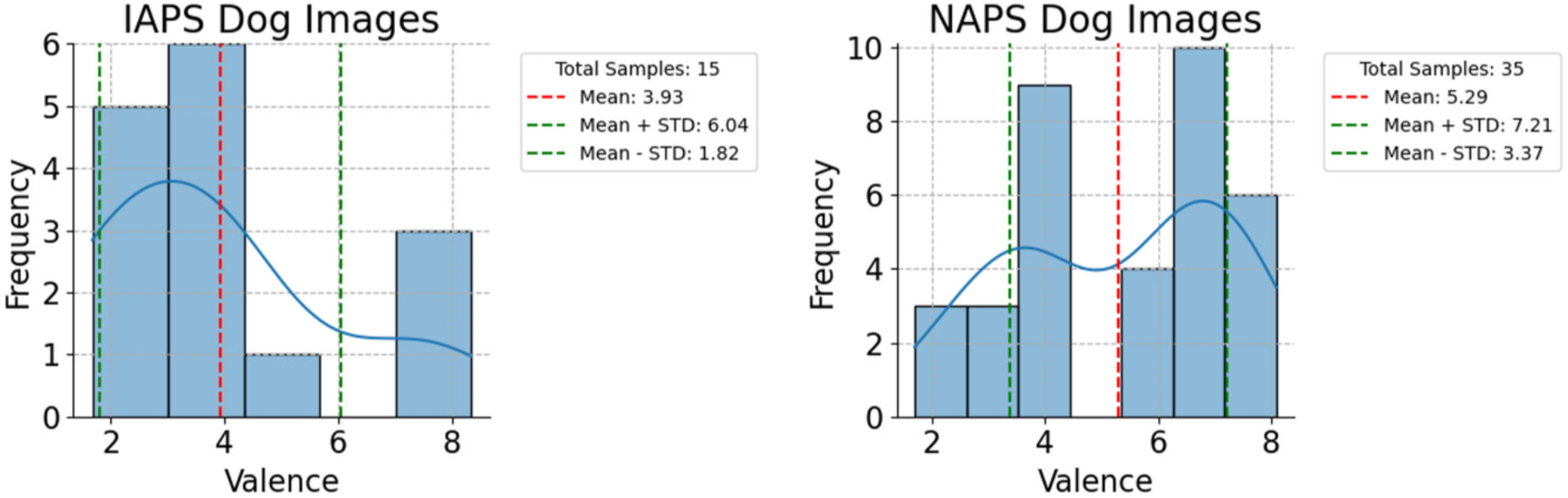
Valence distributions of dog images in dataset IAPS (left) and NAPS (right).

