## Supplemental Figures for "Emergence of Emotion Selectivity in Deep Neural Networks Trained to Recognize Visual Objects"

Eight topics related to the study reported in the main manuscript are addressed in this Supplementary Materials.

**Topic 1. Additional details of model developments, image datasets and methods**

The structure of artificial neurons, the number of images in each dataset, and valence distribution in each dataset are shown in Figure S1. The information flow of applying convolution and ReLU function on an input, the details of how to enhance and lesion artificial neurons, and the datasets and networks used in the study are illustrated in Figure S2.

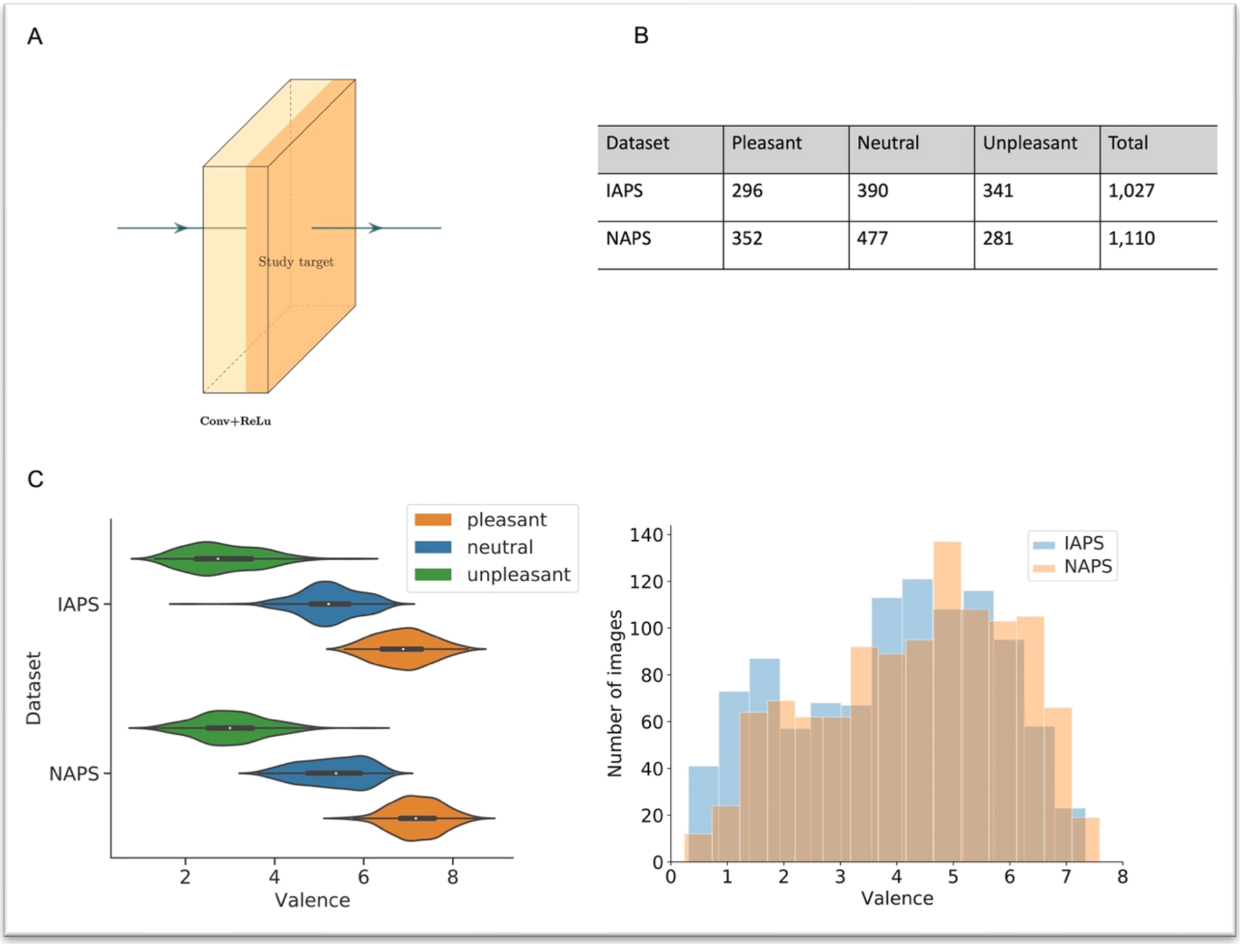


| Dataset | Pleasant | Neutral | Unpleasant | Total |
|---|---|---|---|---|
| IAPS | 296 | 390 | 341 | 1,027 |
| NAPS | 352 | 477 | 281 | 1,110 |

**Figure S1. Model development details and image datasets**. (**A**) Each convolutional layer is followed by one $ReLU$ layer, the output of which reflects the responses of the artificial neurons the convolutional layer. Thus, in this study, the output of the $ReLU$ layer is our study target for understanding the activity of the artificial neurons. (**B**) It shows the number of images of each emotion category in the two datasets used in this study. Two datasets were treated equally for defining emotion-selective neurons and related lesion and attention manipulations. (**C**) It shows how the divided categorial images match the valence score originally rated by human subjects in the two datasets. The C (left) shows the valence score distribution and the boundary score between the pleasant and neutral category: $4.3 \pm 0.5$ and between the neutral and unpleasant category: $6.0 \pm 0.5$. The C (right) shows the number of images per valence score across two datasets. Basically, this figure illustrates the details of the model development and the affective image datasets.

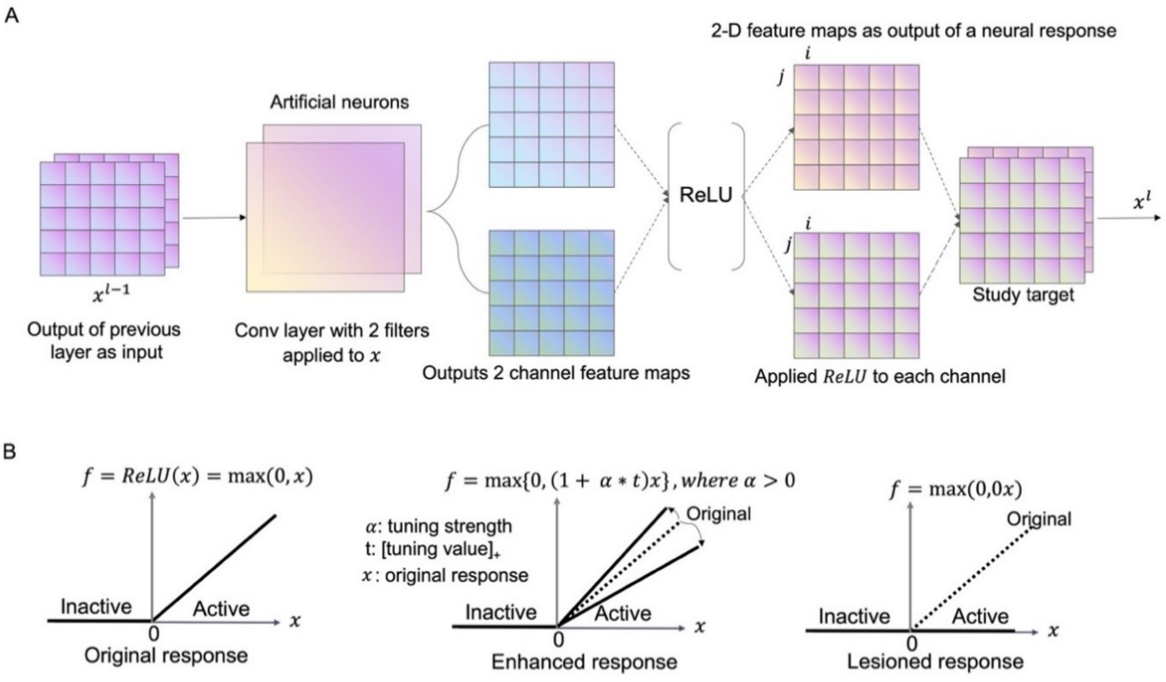


C

The datasets and models used in study

| #emotion categories | Dataset | The usage | Balanced #image per category(e.g., pleasant versus non-pleasant) | Associated model |
|---|---|---|---|---|
| Binary (2 categories) in each dataset | IAPS-pleasant | Tuning study, random sampling study, and lesion study | 296 | IAPS-pl |
| | IAPS-neutral | | 390 | IAPS-ne |
| | IAPS-unpleasant | | 341 | IAPS-un |
| | NAPS-pleasant | | 352 | NAPS-pl |
| | NAPS-neutral | | 477 | NAPS-ne |
| | NAPS-unpleasant | | 281 | NAPS-un |
| - | IAPS | defining selectivity and overlapped neurons for each emotion category | See S. Fig.1B | No need model training |
| | NAPS | | | |

**Figure S2. Further methodological details**. (**A**) It shows that the information flow of applying convolution and $ReLU$ operation to the input $x$ in layer $l$, which represents all the feature maps from the previous layer $l - 1$. In this illustration, $x$ is composed of two channels of feature maps. Two filters (referred to as artificial neurons) in the following convolutional layer are applied to $x$ separately. Each filter convolution results in a channel of a new feature map, which then is passed through an activation function, $ReLU = \max(0, x)$. $ReLU$ function indicates which features after the convolutional operation is activated or inactivated. The active units are valued with positive numbers, and the inactive units are valued with zeros by the $ReLU$. Fundamentally, this figure indicates how an artificial neuron responds to a stimulus and how the response activations are calculated in a CNN. (**B**) The bold black line represents the $ReLU$ response (activation) values, and dash lines in the middle and right sub-figures represent the normal activation value of the $ReLU$. Three cases of activation behavior were investigated empirically. The normal excitation (left) is applying the original $ReLU$; the attention enhanced excitation (middle) is applying a positive weight $\alpha$ to the activation value $x$; the inhabited activation through lesion is setting the activation values to be zeros instead, which performs like a lesion study. (**C**) It summarizes the datasets and the models used in the study. The number of images in the binary model is balanced. The non-* category images were randomly selected from another two categories.

### Topic 2. Additional analysis of the selective index

We examined the distribution and correlation of the selective index (SI) of selective neurons defined on dataset IAPS and NAPS, separately, in each layer and the number of selective neurons across layers by emotion category in IAPS and NAPS. The purpose is to answer the following questions: how many selective neurons there are in the network, how they depend on layers, and how the strength of selectivity depends on layers. The overall selective index is around 0.2 for IAPS and 0.15 for NAPS shown in Figure S3. The correlation between IAPS-defined emotion SI and NAPS-defined SI was computed and the result was shown in Figure S4. The left scatter plot, where neurons from all layers are combined, indicates a positive correlation (Pearson coefficient of 0.30) between IAPS- and NAPS-defined SI. The right plot shows that the correlation between the two SI indices increases as we move deeper into the network. This analysis further supports our claim that emotion selectivity is generalizable across the two datasets.

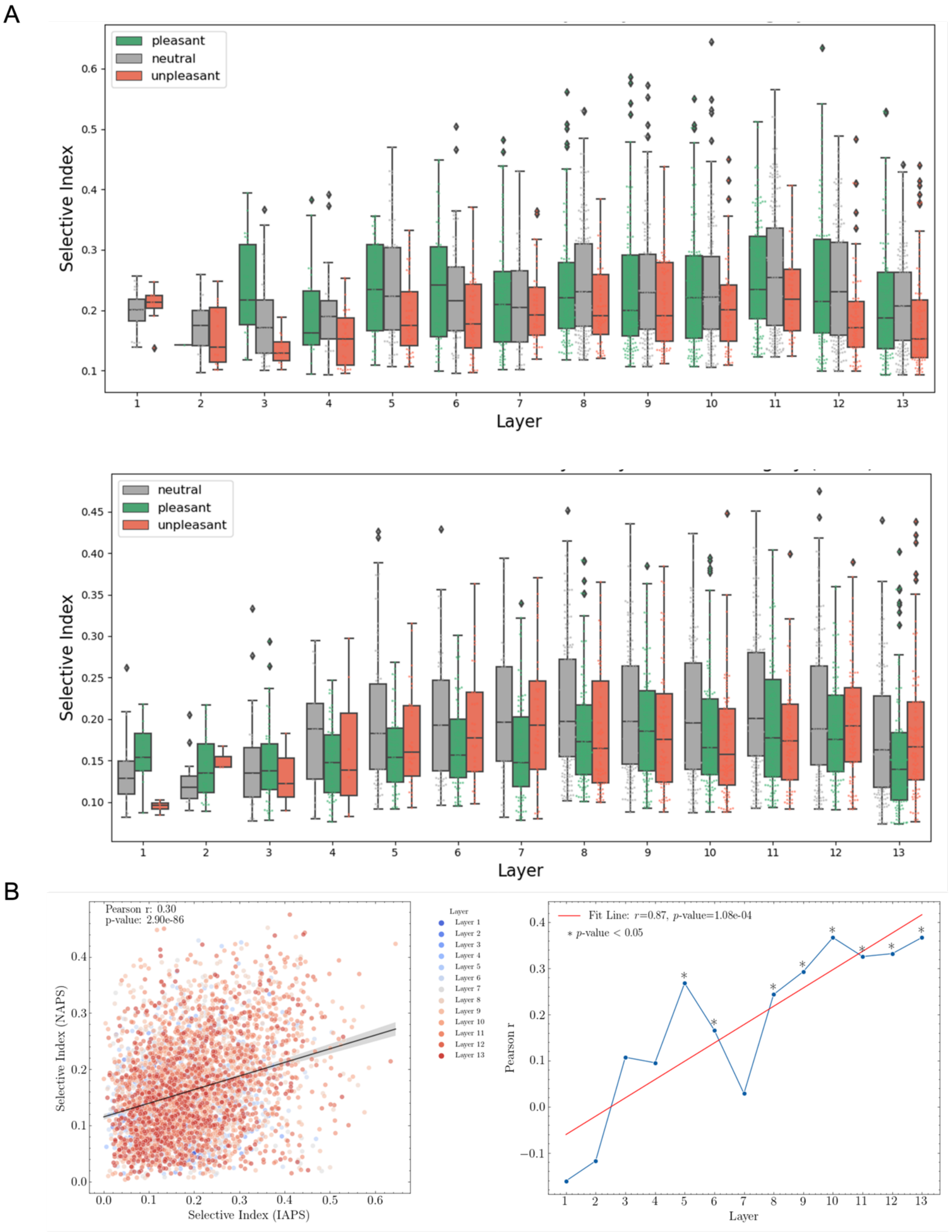


**Figure S3. Additional analysis of selective Index. (A)** Distribution of selective indices across layers by emotion category in dataset IAPS (Top) and NAPS (Bottom). **(B)** The correlation between IAPS-defined SI and NAPS-defined SI. (Left) Neurons from all layers are combined. (Right) The layer-wise correlation was plotted.

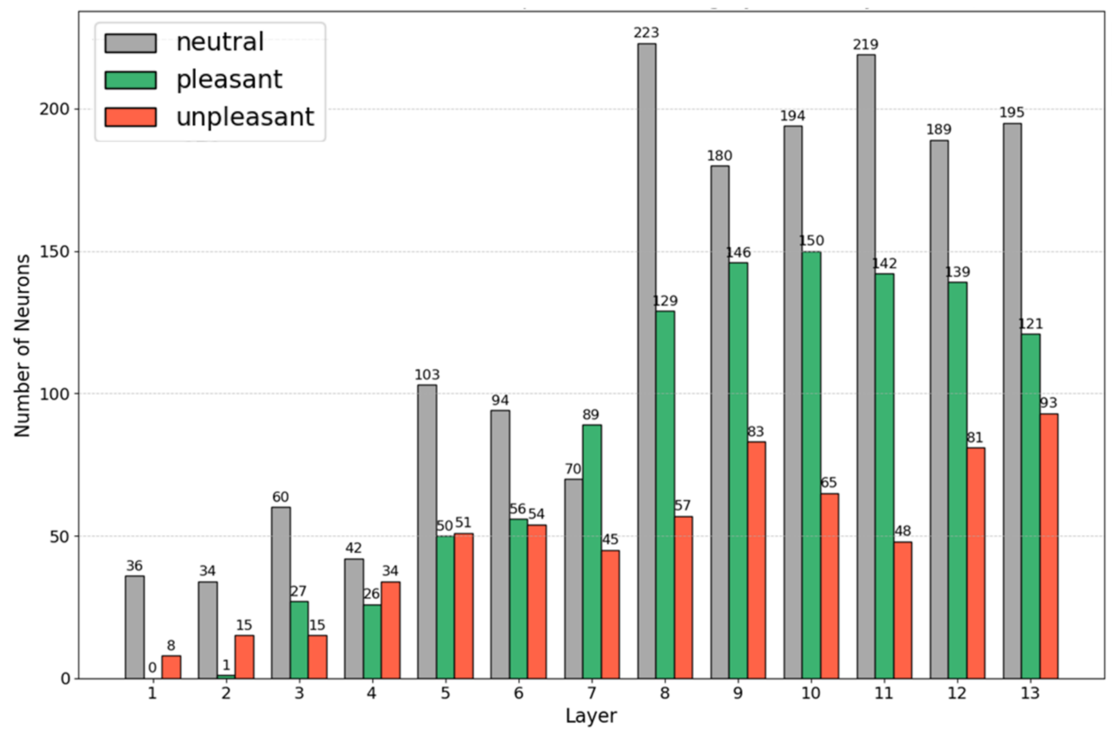


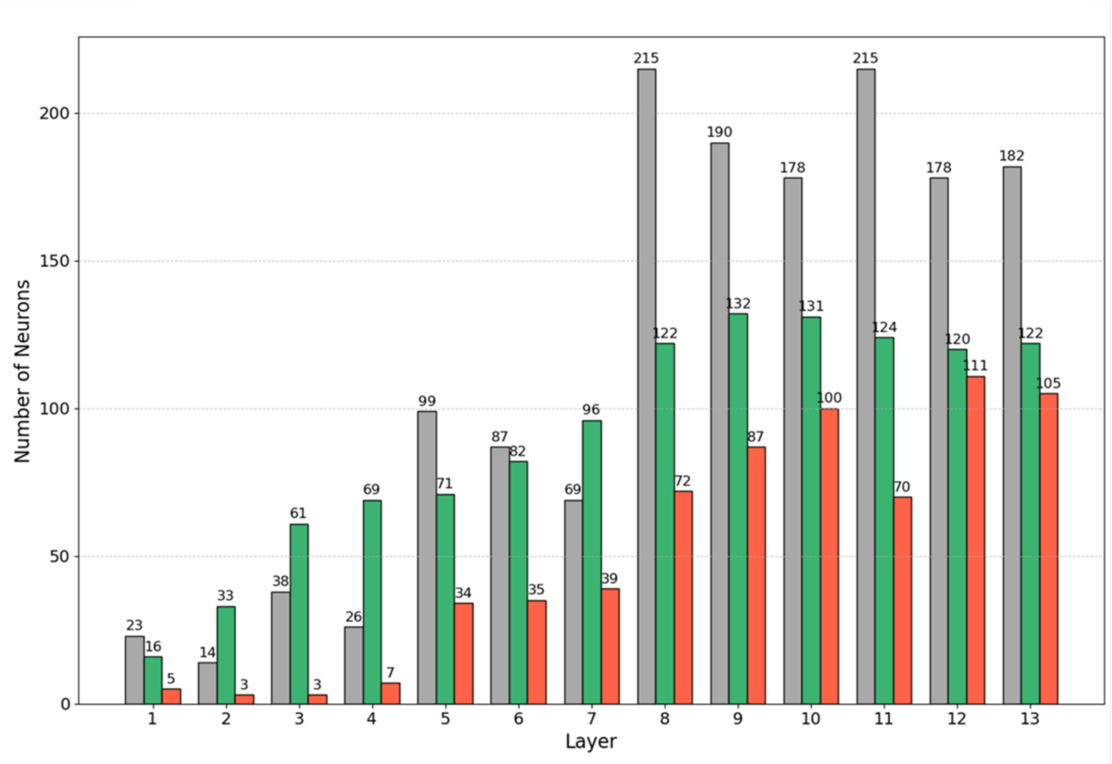


**Figure S4. Number of selective neurons across layers by emotion category in dataset IAPS (Top) and NAPS (Bottom).**

### Topic 3. Generalizability of emotion selectivity

We examined the functional generalization of selective neurons defined on IAPS and NAPS, separately, in Figure S5. The purpose is to further verify whether the emotion selectivity defined on one dataset can be functionally generalized to another dataset. The result is consistent with other results obtained by enchaining on selective neurons with their selective index defined on the same dataset (either IAPS or NPAS). It further supports our claim that emotion selectivity shares a functional property between the two datasets.

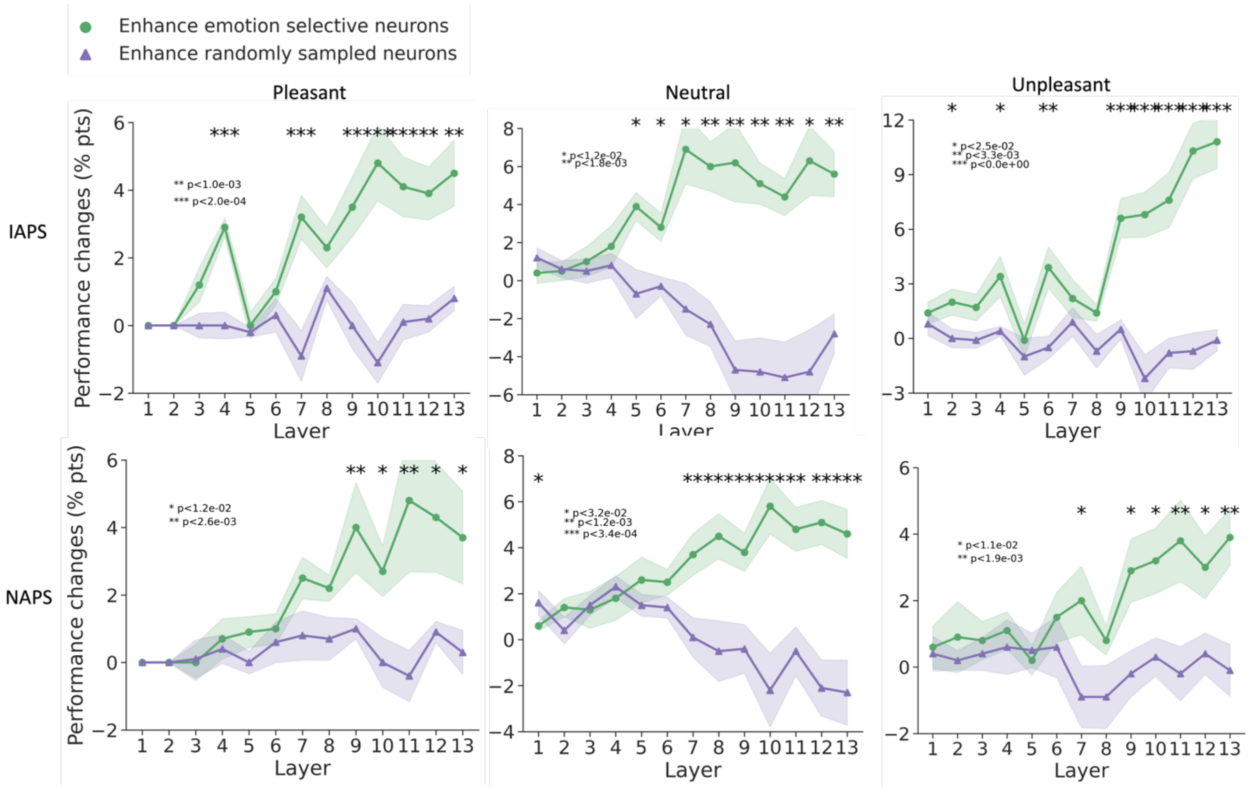


**Figure S5. Functional generalization analysis.** (**Top**) We analyzed the enhancement of emotion-selective neurons (defined post-threshold) versus random neurons in each VGG-16 layer trained on IAPS. The selective index was derived from NAPS and tested on IAPS. (**Bottom**) We analyzed the enhancement of emotion-selective neurons (defined post-threshold) versus random neurons in each VGG-16 layer trained on NAPS. The selective index was derived from IAPS and tested on NAPS.

### Topic 4. Result replication in AlexNet

We replicated the results in another network, AlexNet, shown in Figure S6, Figure S7, and Figure S8. The purpose is to demonstrate that the emergence of emotion selectivity is not an idiosyncratic property of a specific deep neural network. We summarized the parallel results produced with VGG-16 and AlexNet in Table S1 and Table S2.

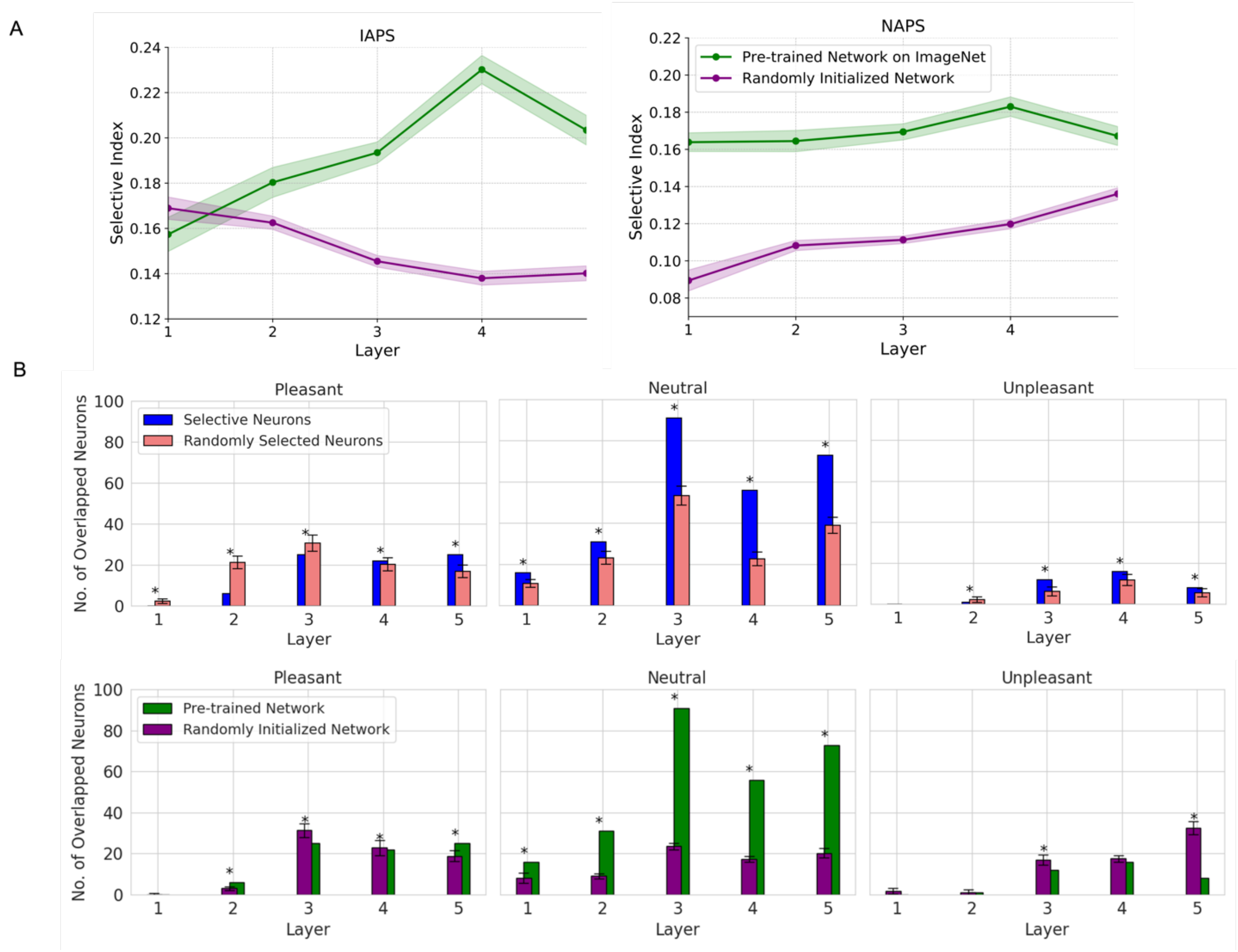


**Figure S6. Selective Index Quality (A) and Generalizability in AlexNet (B)** of emotion selectivity across two datasets**.** The comparison of number of overlapped neurons derived from selective neurons and randomly selected neurons is plotted (B-top). The one of number of overlapped neurons derived from pre-trained AlexNet on ImageNet and initialized AlexNet network with random weights (B-bottom). The goal of this comparison is to demonstrate the significance of learned features from ImageNet in developing neuron selectivity. However, merely counting the overlapping neurons might not be adequate; we should also take into account the selectivity index quality. This is particularly important when the total number of neurons in a layer is small.

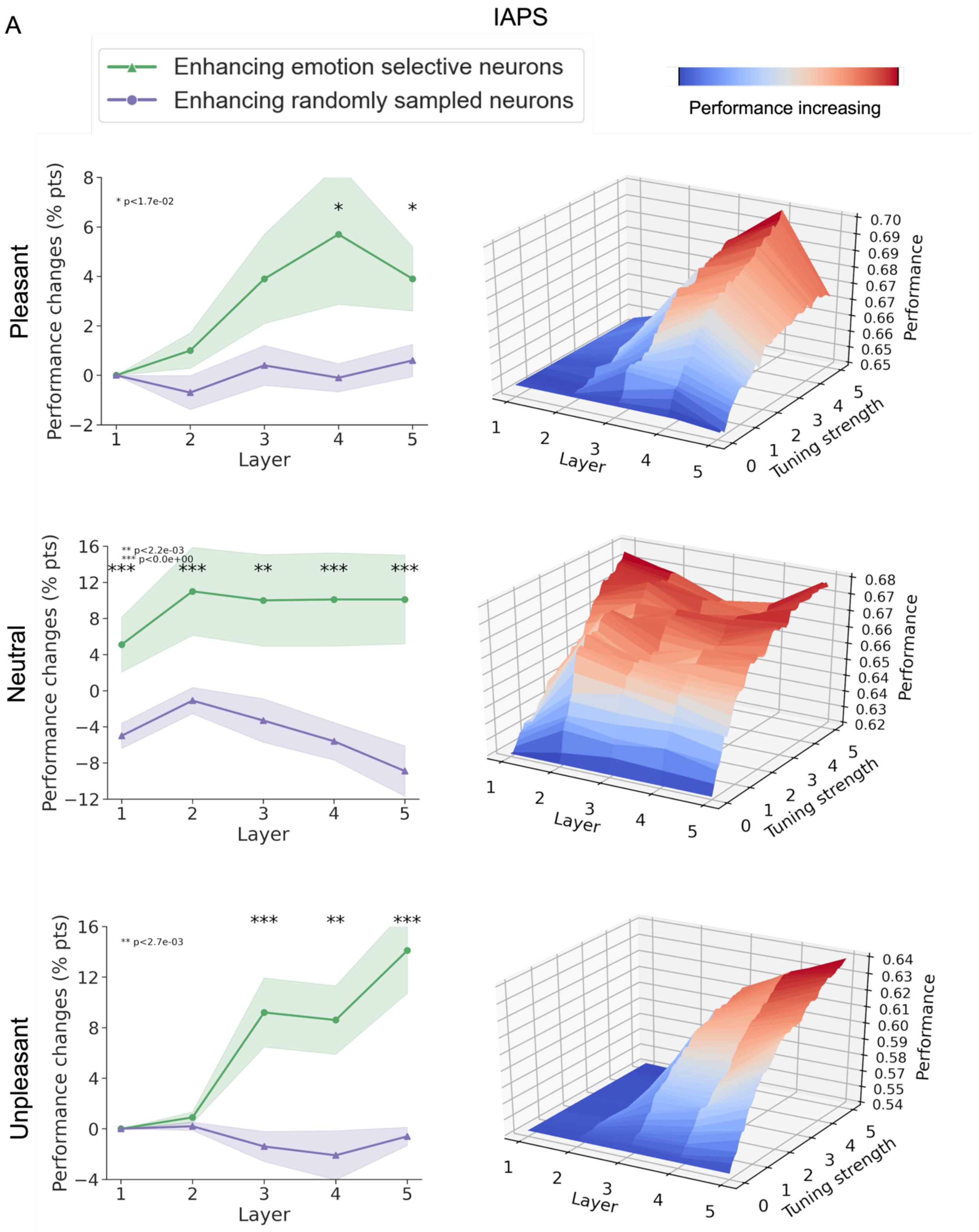


**Figure S7. Effects of attention-enhancing emotion-selective neurons and randomly selected neurons in AlexNet.** (**A**) IAPS dataset. (**B**) NAPS dataset.

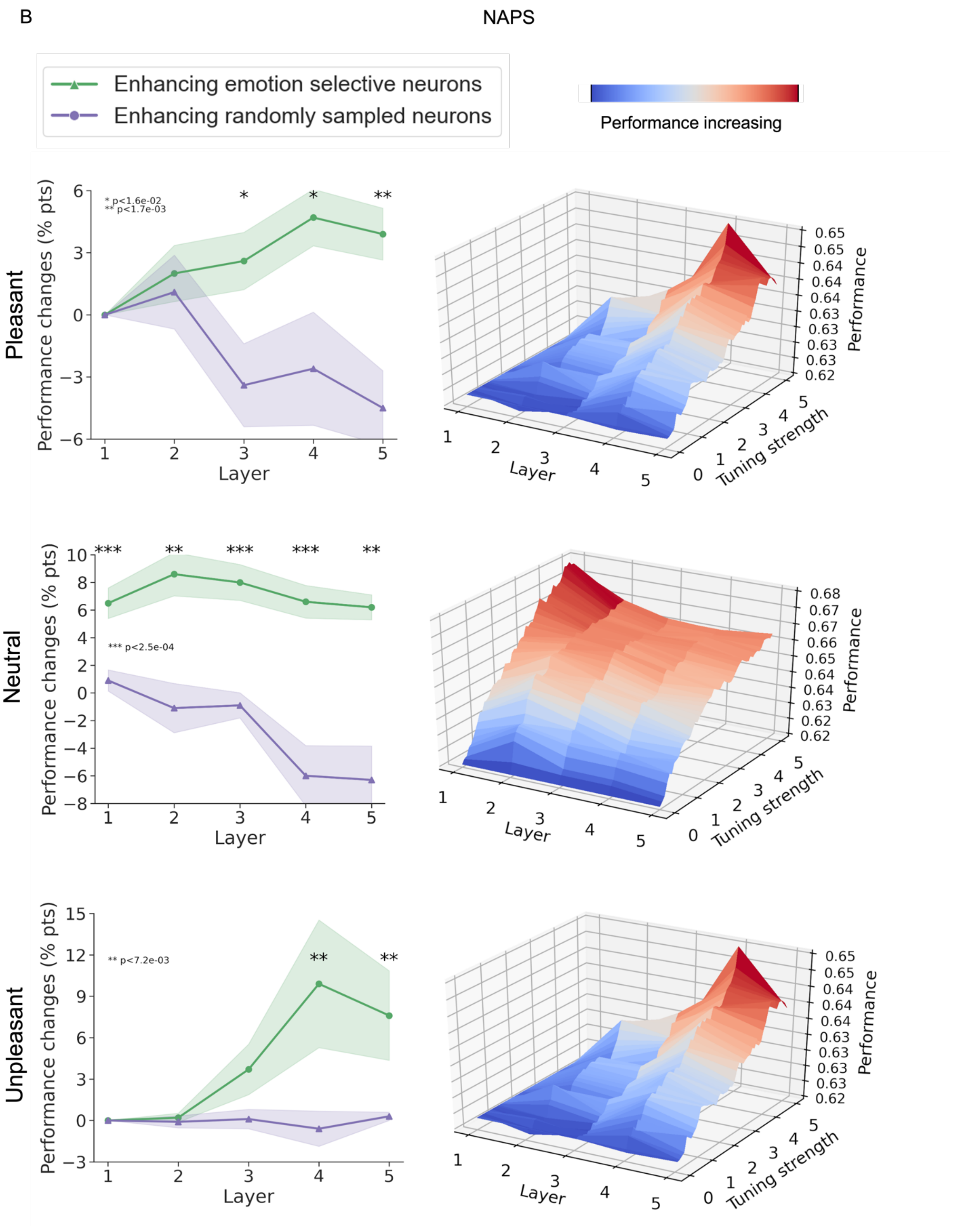


**Figure S7. Continued**

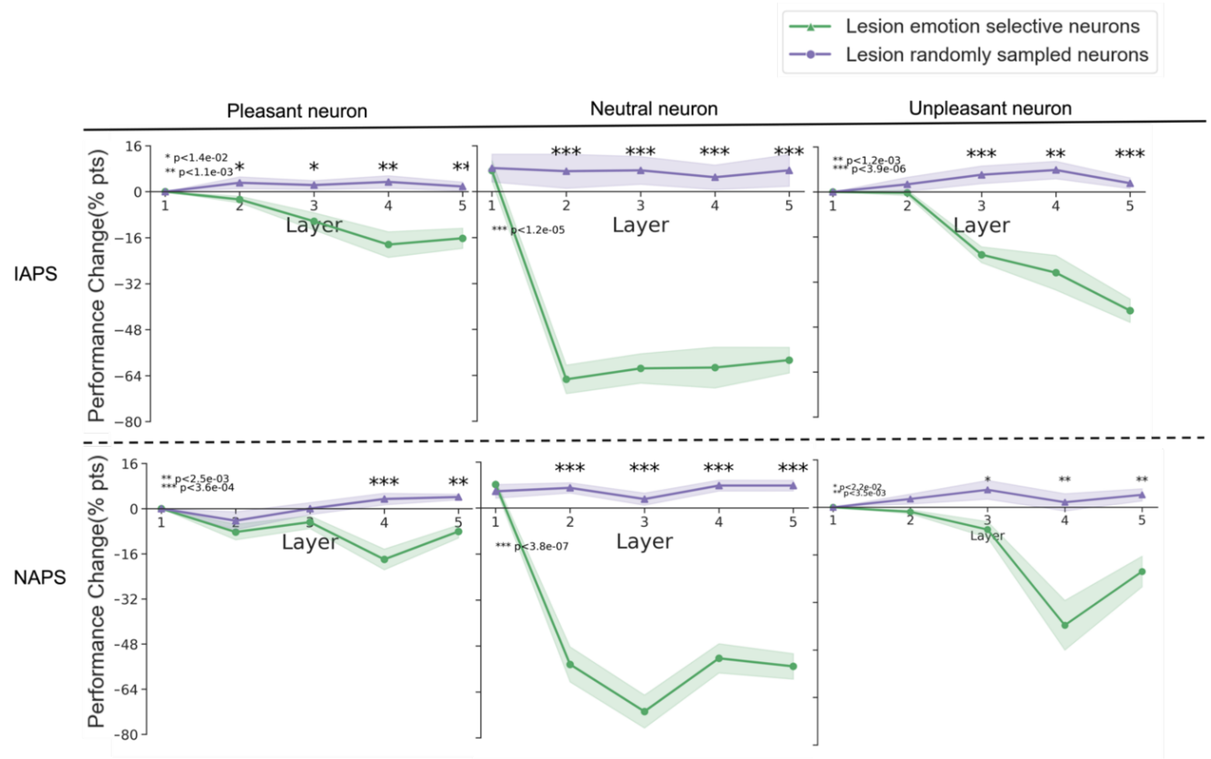


**Figure S8. Effects of lesioning emotion-selective neurons and randomly selected neurons in AlexNet.** (**A**) IAPS dataset. (**B**) NAPS dataset**.**

**Table S1. Results comparison between VGG-16 and AlexNet.**

| **Description** | **VGG-16** | **AlexNet** |
|---|---|---|
| Selective Index Quality | Fig.3A | Fig.S3 |
| Number of overlapped neurons | Fig.3B | Fig.S4 |
| Enhance emotion- selective neurons | Fig.4 | Fig.S7 |
| Lesion emotion-selective neurons | Fig.5 | Fig.S8 |

**Table S2 Original and Enhanced Performance (F1-score) in VGG-16 and AlexNet.**

| Network | Dataset | Emotion to Recognize | Original Performance | Enhanced Performance | Enh. Improvement (%) | Lesioned Performance | Les. Decreased (%) |
|---|---|---|---|---|---|---|---|
| VGG-16 | IAPS | Pleasant | 0.70 | 0.73 | 4.29% | 0.56 | 20% |
| | | Neutral | 0.63 | 0.69 | 9.52% | 0.26 | 58% |
| | | Unpleasant | 0.62 | 0.69 | 11.29% | 0.13 | 80% |
| | NAPS | Pleasant | 0.70 | 0.72 | 2.86% | 0.49 | 31% |
| | | Neutral | 0.63 | 0.67 | 6.35% | 0.25 | 61% |
| | | Unpleasant | 0.67 | 0.71 | 5.97% | 0.41 | 39% |
| AlexNet | IAPS | Pleasant | 0.65 | 0.70 | 7.69% | 0.55 | 16% |
| | | Neutral | 0.62 | 0.68 | 9.68% | 0.22 | 64% |
| | | Unpleasant | 0.54 | 0.64 | 18.52% | 0.37 | 32% |
| | NAPS | Pleasant | 0.62 | 0.65 | 4.84% | 0.52 | 16% |
| | | Neutral | 0.62 | 0.68 | 9.68% | 0.22 | 65% |
| | | Unpleasant | 0.62 | 0.65 | 4.84% | 0.40 | 35% |

### Topic 5. Low-level features as possible confounding factors

low-level features were extracted from the images by using GIST algorithm. Pairwise emotion decoding was performed using (see Figure S9) using SVM. The mean accuracy for both IAPS and NAPS datasets approximates the chance level, suggesting that low-level GIST features are insufficient for decoding emotion categories from images.

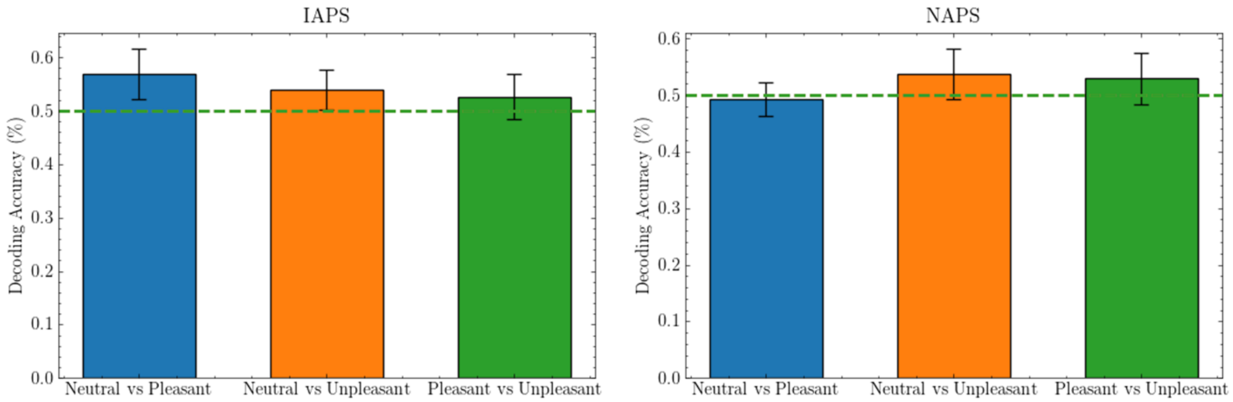


**Figure S9. Pairwise decoding results using low-level features (GIST).** Dash line indicates the chance level performance (50%). The dashed lines indicate chance-level performance and bars representing the average accuracy across 10 iterations of 5-fold cross-validation.

### Topic 6. Faces as possible confounding factors

The percentages of images involving faces in top 100 images (ranking based on neurons' activation to each image) that evoked the strongest response of selective neurons for each emotion category (see Figure S10) are: 16%, 62%, and 30% for pleasant, neutral, and unpleasant, separately. The analysis demonstrates that development of emotion selectivity in these neurons is unlikely affected by potential facial encoding that might arise during the training of the network on ImageNet.

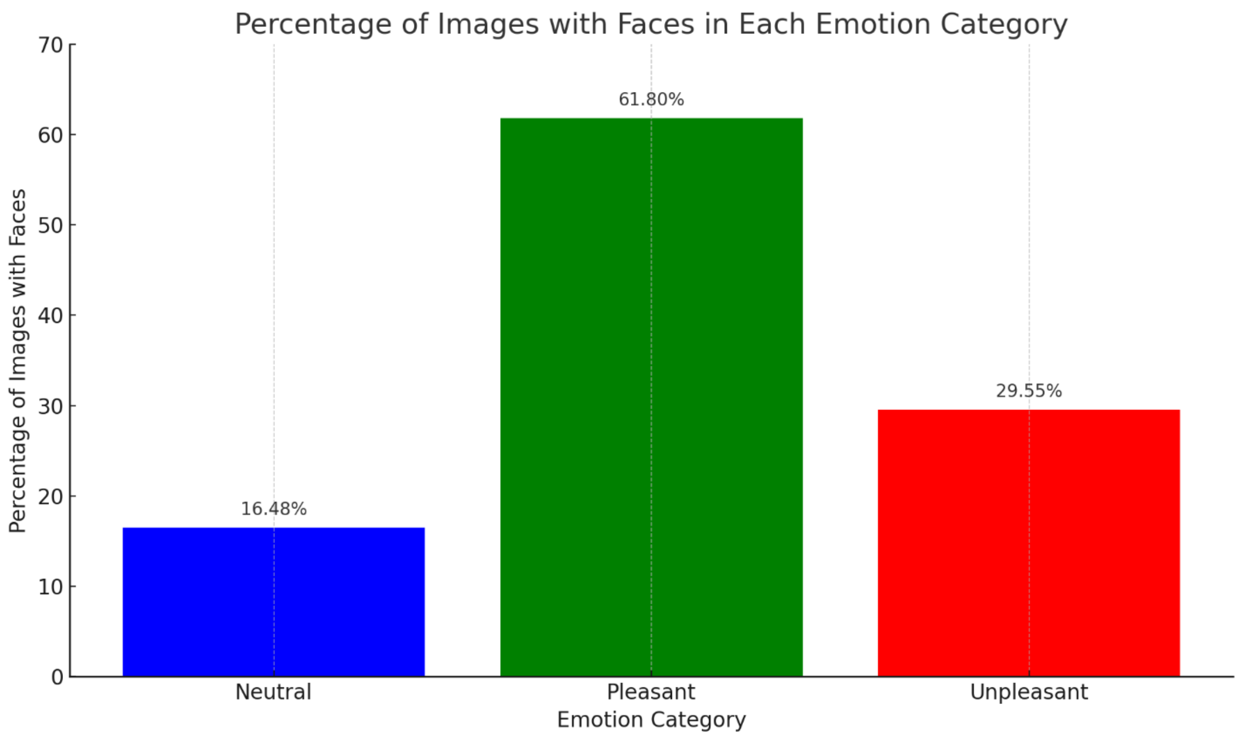


**Figure S10. Number of images involving faces in top 100 images that evoked the strongest response of emotion-selective neurons.**

### Topic 7. Animacy as possible confounding factors

Table S3 shows the mean valence and arousal of the top 100 images across selective neuron categories. The purpose of this analysis is to estimate how much valence and arousal relevant to the images evoked by the selective neurons are captured. The result shows the mean valence: 6.770, 5.180, and 2.898 and mean arousal: 5.055, 3.970, and 5.816 for top images that evoked the strongest responses in neurons selective for pleasant, neutral, and unpleasant emotion. More importantly, these images appear to contain both animate and inanimate content, suggesting that animacy might not be a confounding factor.

**Table S3 Valence and Arousal of Top 100 Images Across Selective Neuron Categories.** Note: Image ranking is based on neurons' activation to each image.

| Neuron Selectivity | Mean Valence | Interpretation | Mean Arousal | Interpretation |
|---|---|---|---|---|
| Pleasant | 6.770 | Images evoke relatively positive or pleasant emotions. | 5.055 | Emotions of moderate intensity. |
| Neutral | 5.180 | Emotions neither particularly positive nor negative. | 3.970 | More subdued or calm emotions. |
| Unpleasant | 2.898 | Images evoke negative or unpleasant emotions. | 5.816 | Intense negative emotions (e.g., fear, distress). |

**Topic 8. Effects of emotion, object category, and their interaction on neuronal responses**

As illustrated in Figures S11-S12, we examined, using a Two-Way ANOVA analyses, the impact of image emotion, image category, and their interaction on neuronal response. Object categories were identified based on the descriptions in the original datasets (refer to Figure S11A and S12A). Our findings reveal that the emotion category markedly affects neuronal activity in layers subsequent to the fifth (refer to Figure S11B-top and S12B-top), and the influence of the object category is increasing with layer depth but not significant. The interaction is significant in some deeper layers (refer to Figure S11B-bottom and S12B-bottom). It should be noted that this analysis should be viewed as preliminary, because the number of images in each object category is rather small, which may impact the analysis adversely. In addition, the selection of the images is also dataset-specific. For example, in the IAPS dataset, 15 images of dogs predominantly express negative emotions, whereas in the NAPS dataset, 35 images of dogs represent a mix of negative and positive emotions (see Figure S13). This variance indicates the necessity for additional studies to comprehensively understand the interaction between image emotion and category.

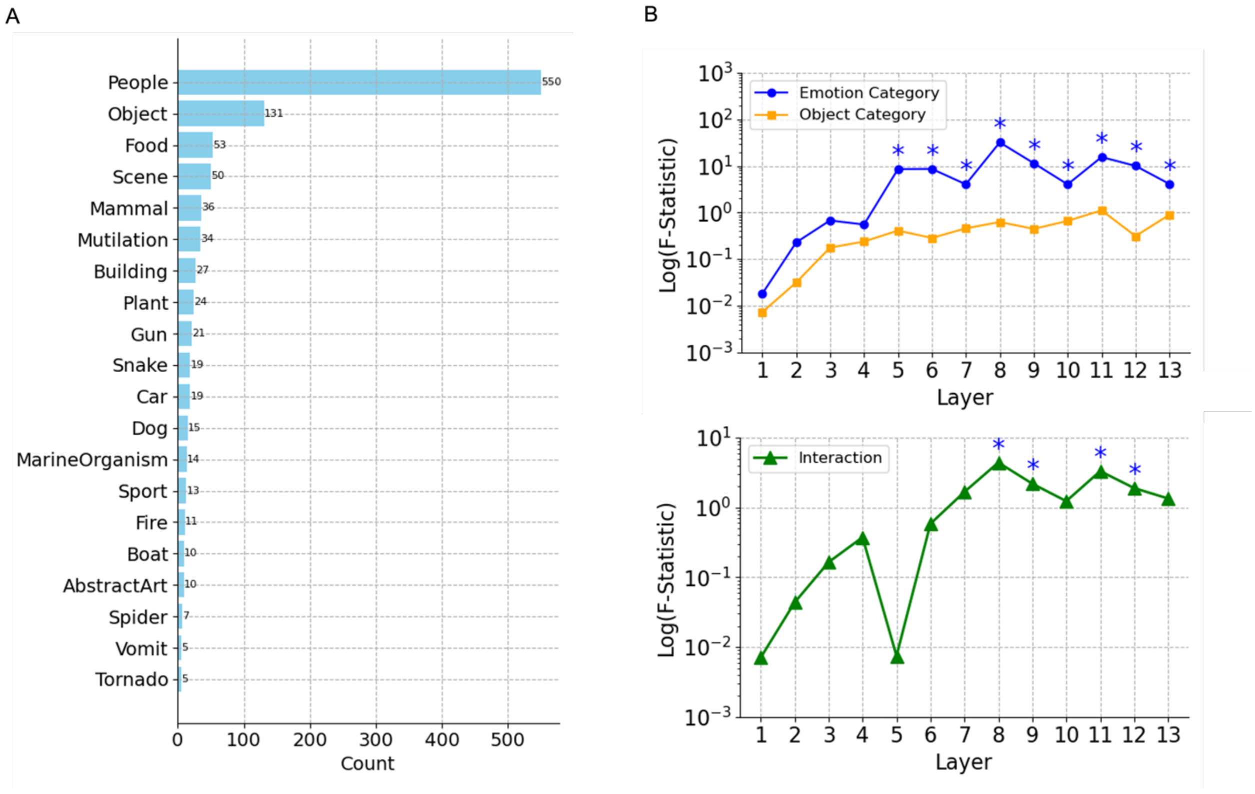


**Figure S11. Effects of emotion and object category on filter activity using *IAPS* images. A.** The number of images in each of the top 20 object categories. **B. (top)** The F-statistic (log scale) of the effect of emotion and object category on filter activations across layers of the VGG-16 neural network. The statistics are obtained from a Two-Way ANOVA test, where the dependent variable is the filter activity in response to images. The plot reveals how each factor impacts the filter responses and how this influence changes from the input to deeper layers of the network; (**bottom**) The F-statistic (log scale) of the interaction between emotion dependent filter activation and object category dependent filter activation. The statistics are obtained from a Two-Way ANOVA test. * *indicates the influence is statistically significant.*

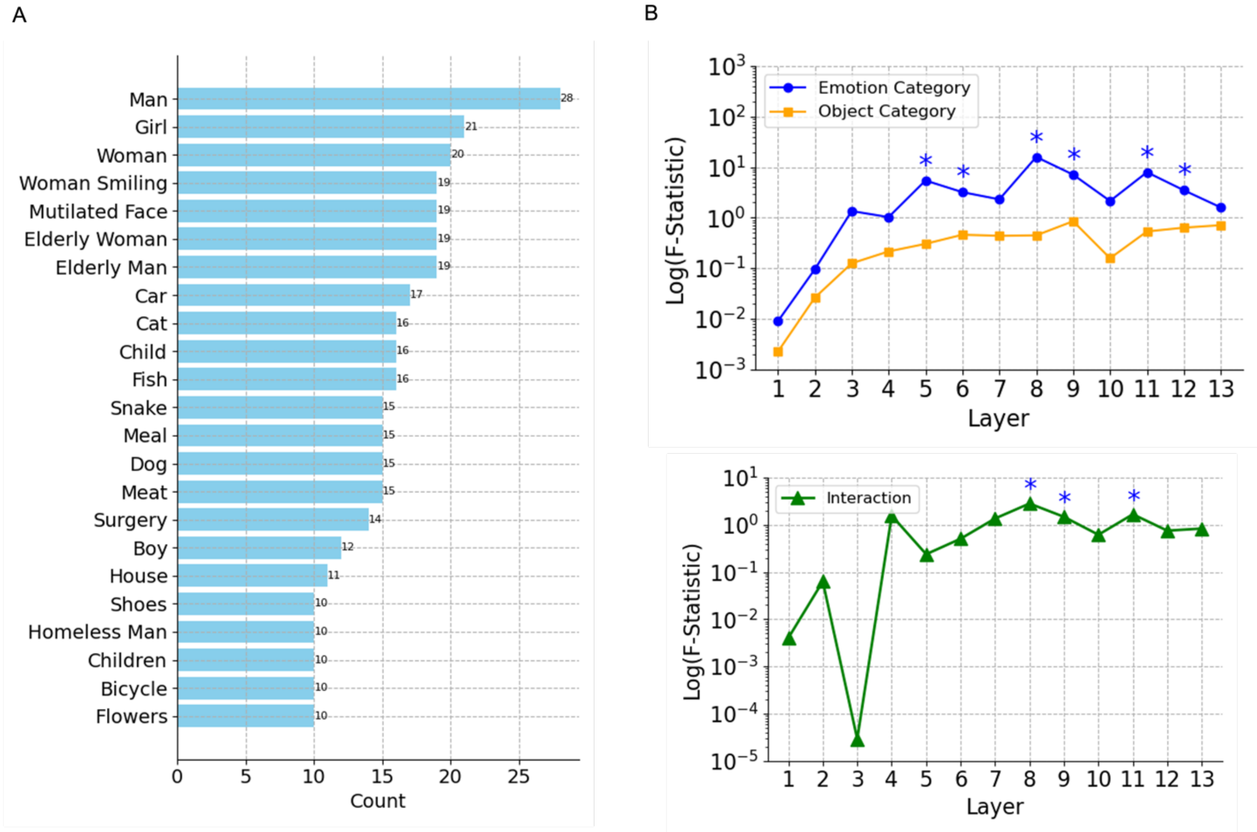


**Figure S12**. **Effects of emotion and object category on filter activity using *NAPS* images**. **A.** The number of images in each of the top 23 object categories. **B. (top)** The F-statistic (log scale) of the effect of emotion and object category in filter activations across layers of the VGG-16 neural network. The statistics are obtained from a Two-Way ANOVA test, where the dependent variable is the filter activity in response to images. The plot reveals how each factor impacts the filter responses and how this influence changes from the input to deeper layers of the network; (**bottom**) The F-statistic (log scale) of interaction between emotion and object category. The statistics are obtained from a Two-Way ANOVA test. * *indicates the influence is statistically significant.*

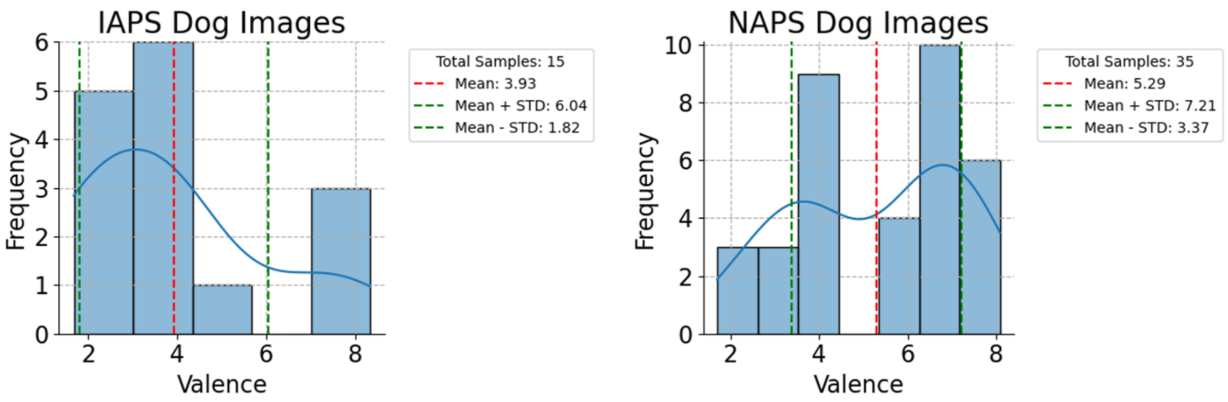


**Figure S13. Valence distributions of dog images in dataset IAPS (left) and NAPS (right).**
